# Age-related increase in matrix stiffness downregulates α-Klotho in chondrocytes and induces cartilage degeneration

**DOI:** 10.1101/2021.03.13.434679

**Authors:** Hirotaka Iijima, Gabrielle Gilmer, Kai Wang, Allison Bean, Yuchen He, Hang Lin, Christopher Evans, Fabrisia Ambrosio

**Author notes:** **Corresponding author:** Fabrisia Ambrosio, PhD, MPT, Suite 308, Bridgeside Point Building II, 450 Technology Drive, Pittsburgh, PA 15219. **Co-corresponding author:** Hirotaka Iijima, PhD, PT, Suite 308, Bridgeside Point Building II, 450 Technology Drive, Pittsburgh, PA 15219.

## Abstract

Enhanced mechanistic insight into age-related knee osteoarthritis (KOA) is an essential step to promote successful translation of animal research to bedside interventions. To this end, the goal of these studies was to interrogate molecular mechanisms driving age-related KOA in a mouse model and correspond findings to human knee cartilage. Unbiased mass spectrometry proteomics of cartilage tissue revealed PI3K/Akt signaling was the predominant pathway disrupted over time in male, but not female, mice. This finding was consistent with a significantly accelerated KOA progression in males when compared to female counterparts. In probing for upstream regulators of these age-dependent alterations, we found that α-Klotho, a suppressor of PI3K/Akt signaling and potent longevity protein, significantly decreased with aging in both mouse and human knee cartilage. Upstream of these alterations, we found that age-related increases in matrix stiffness initiated a cascade of altered nuclear morphology and downregulated α-Klotho expression, ultimately impairing chondrocyte health. Conversely, reducing matrix stiffness increased α-Klotho expression in chondrocytes, thus enhancing their chondrogencity and cartilage integrity. Collectively, our findings establish a novel mechanistic link between age-related alterations in ECM biophysical properties and regulation of cartilage health by α-Klotho.

## INTRODUCTION

All cells in the human body are subject to mechanical influences. This is particularly true of articular cartilage, given that its primary role is to transmit forces to the underlying bone and decrease friction in the joint. However, these important functions are disrupted by the progressive cartilage deterioration that occurs with aging. In 1742, British anatomist William Hunter wrote “…*articular cartilage, when it is injured or possesses pathology, does not heal*”(*1*). In most cases, cartilage damage eventually progresses to osteoarthritis (OA), which affects approximately 32.5 million Americans(*2*). Although we have known of the poor healing capacity of cartilage for over 250 years, this limited capacity is both poorly understood and inadequately treated in the clinic.

The lack of successful non-surgical treatment for OA is attributed, in part, to 1) gaps in our current understanding of whether and how pre-clinical OA models recapitulate human disease and 2) an incomplete understanding of the complete trajectory of molecular mechanisms driving disease development. This is particularly true of knee OA (KOA). In most cases, there is no clear inciting event in KOA, and the single greatest predictive factor of onset is age(*3*). Post-menopausal women are also disproportionately diagnosed with KOA(*4*). Despite this prevalence, phenotypic alterations in knee cartilage over time and according to sex have not been thoroughly described. To date, the majority of animal studies have utilized a post-traumatic model of OA (PTOA) in young male mice(*5*). This post-traumatic model displays a distinct trajectory of molecular perturbations in response to gene modification when compared to age-associated KOA(*6*). Thus, it is not surprising that the majority of pre-clinical findings in OA have failed to translate to treatment for a population consisting largely of aged individuals with no joint trauma.

In this study, we thoroughly characterize the trajectory of KOA in mice across the lifespan and according to sex, and we compare it to our current understanding of clinical KOA. As a part of this characterization, we performed mass spectrometry proteomics. Protein-level changes converged on the PI3K/Akt signaling pathway as significantly associated with age-related cartilage degeneration. Using loss-of-function paradigms, we then identified α-Klotho, a longevity protein and suppressor of PI3K/Akt signaling, as a novel regulator of chondrocyte health. In order to better understand the mechanisms underlying a loss of α-Klotho over time, we performed a series of loss-of-function paradigms *in vitro* and *in vivo* and found that age-related biophysical ECM alterations drive decreased α-Klotho expression and phenocopy clinical features of KOA.

## RESULTS

### Aging induces accelerated cartilage degeneration in males relative to female counterparts

To characterize the natural trajectory of age-associated KOA in mice, we evaluated cartilage integrity in three age groups of male and female C57/BL6 mice: young (4-6 months), middle-aged (10-12 months), and aged (18-24 months) (**Figure 1A**). These age groups correspond to 20-30, 38-47, and 56-69 years of age in humans, respectively(*7*). This study focused on medial tibial cartilage given this is the region most typically affected in humans(*8*).

**Figure 1.**
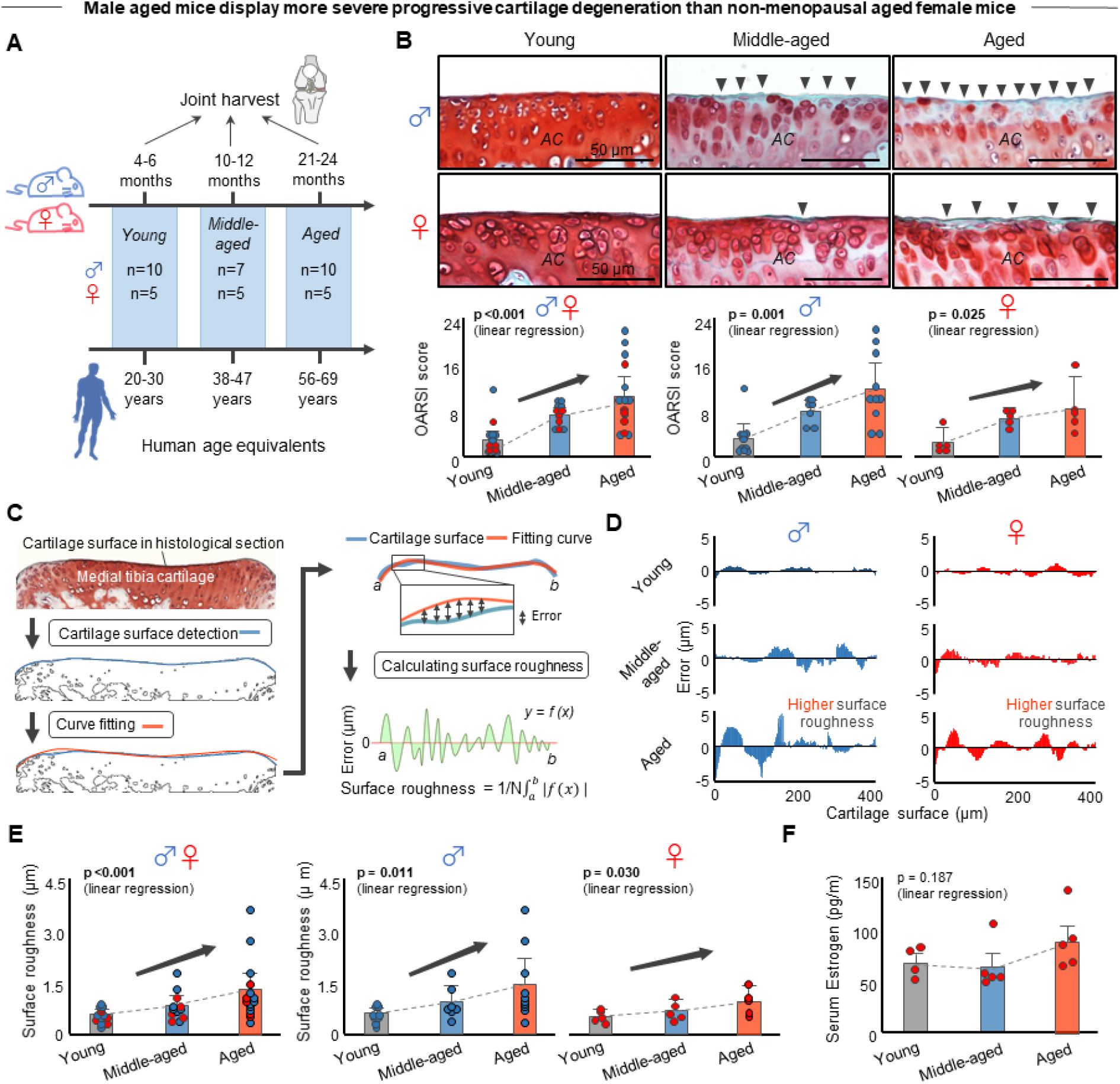
Aging induced progression cartilage degeneration and surface roughness in a sex-dependent manner. **A.** Schematic showing the experimental protocol. Human age equivalent is provided. **B.** Aging induces progressive cartilage degeneration in murine medial tibial plateaus in a sex-dependent manner. Representative histological sections stained with Safranin-O/Fast Green are provided in each group. Black arrow heads indicate loss of cartilage matrix. OARSI score (0-24 points; higher value indicates more severe cartilage degeneration) assessed by blinded assessor is provided. **C.** Computational analysis for calculation of cartilage surface roughness. Surface roughness was calculated as the deviation between actual cartilage surface (blue solid line) and fitted curve applied to the cartilage surface (red solid line). **D.** Representative error value between fitting curve and cartilage surface. **E.** Aging induces progressive surface roughness in murine medial tibial plateaus in a sex-dependent manner. **F.** Similar serum Estrogen level across three groups in female mice (n = 4 in young, n = 5 in middle-aged and aged). Statistical analysis was performed using linear regression analysis (**B**, **E**, **F**). Data are presented as means ± 95% confidence intervals.

Histological findings confirmed progressive cartilage degeneration beginning at middle-age. These findings are in line with clinical reports of structural abnormalities that manifest in human cartilage at middle-age(*9, 10*) (**Figure 1B**). To further support the progressive cartilage degeneration, we quantified cartilage surface roughness by assessing the deviation of the cartilage surface from a fitted curve (**Figure 1C**). As expected, surface roughness significantly increased with aging (**Figure 1D, E**). Moreover, the magnitude of cartilage degeneration and increased surface roughness was greater in male mice than in female mice(**Figure1D,E**). It is unclear why female mice appear to display relative protection against KOA. However, it is worth noting that serum estrogen levels were unchanged across the three female age groups, indicating that aged female mice do not experience the same menopausal phenotypic changes seen in humans (**Figure 1F**). Previous studies have demonstrated that approximately 65% of aged female mice spontaneously transition to a polyfollicular anovulatory state, with estrogen and progesterone profiles resembling that of premenopausal women(*11*). This is especially important since post-menopausal women typically present with more severe KOA than men(*12*).

### Mass spectrometry proteomics reveal age-related enrichment of PI3K-Akt signaling pathway in male, but not female, mice

We next performed mass spectrometry proteomics to explore potential signaling pathways associated with sex- and age-dependent cartilage degeneration. Articular cartilage was collected from young, middle-aged, and aged male and female mice (n=5/age/sex; **Figure 2A**). From these samples, we identified 44,689 peptides associated with 6,694 unique proteins (**Figure 2A**). Data are available via ProteomeXchange with identifier PXD024062 (see **Methods** for details).

**Figure 2.**
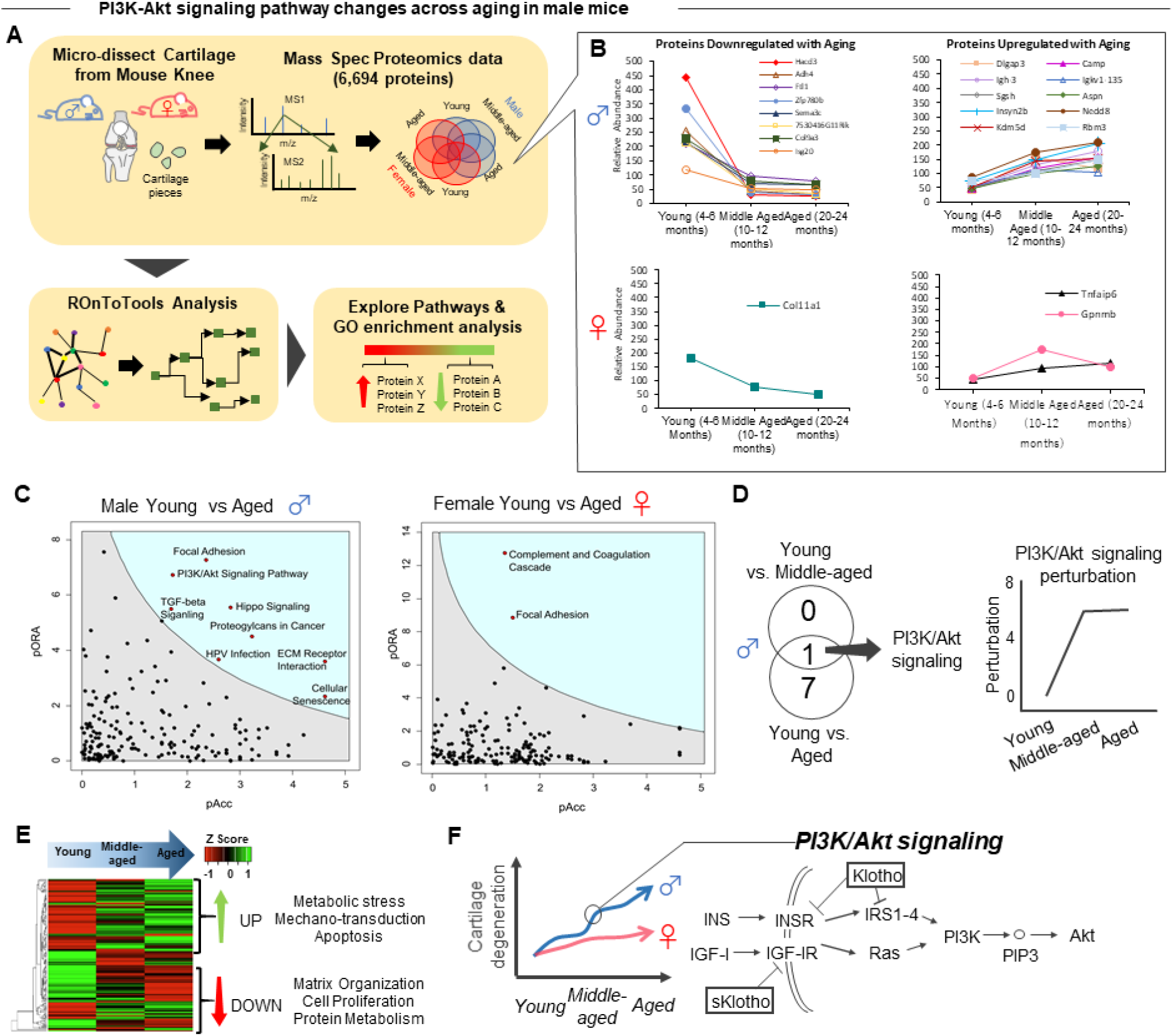
Mass spectrometry proteomics reveal age-enrichment of PI3K-Akt signaling pathway in male mice but not female mice. **A.** Schematic showing the experimental protocol. Knee cartilage was micro-dissected from young, middle-aged, and aged male and female mice (n=5/sex/age). **B.** Individual proteins up-and downregulated with aging for male and female mice. **C.** Pathway analysis for young vs. aged in male and female mice. **D.** Comparison between young vs. aged and young vs. middle-aged analyses in male mice, and display of perturbation calculations. **E.** Heat map of detected proteins that are involved with PI3K/Akt signaling. **F.** Schematic correlating Figure 1 findings of cartilage degeneration to pathway involving PI3K/Akt signaling and linking to α-Klotho.

Nine proteins were significantly different between young and middle-aged female samples, and thirty-eight proteins were significantly different when comparing young and aged female cartilage (**Figure S1, S2**). Of these, only three proteins were significantly different in both the young vs. middle-aged and young vs. aged comparisons (**Figure 2B**). A similar comparison was performed between sexes (male vs. female) across all three age groups and is available in **Figures S3-S5**. The most prominent changes occurred in immune-related proteins, suggesting that differences in immune function may play a role in the observed sex differences.

Age-associated trends in male mice were notably different when compared to female counterparts. Twenty-three proteins were significantly different between young and middle-aged male mice, and fifty-six proteins were significantly different between young and aged male mice (**Figure S6, S7**). Of these, 20 differentially expressed proteins were common to both the young vs. middle-aged and young vs. aged male comparisons. Compared to female mice, male mice demonstrate far more extensive changes in protein content with aging, suggesting distinct sex-related differences in cartilage-aging phenotype. Interestingly, the greatest magnitude of protein changes was observed between the young and middle-aged samples (**Figure 2B**).

Next, we performed Kyoto Encyclopedia of Genes and Genomes (KEGG) enrichment analyses to gain a holistic understanding of the protein network level changes in knee cartilage with aging(*13*). As suggested by the individual protein data, few pathways changed over time in female mice, with only one significantly enriched pathway between young and middle-aged samples and only two pathways differentially expressed between young and aged samples (**Figure 2C & S8, S9**). Of these, *Coagulation and Complement Cascade* changed in both the young vs. middle-aged and young vs. aged comparisons. The complement cascade is activated in OA through mechanical, catabolic, and inflammatory mediators(*14*), and studies have shown complement is elevated in human joints with OA(*15*). A similar pathway analysis was performed comparing males vs. females across the three age groups and is available in **Figures S10-S12**. Similar to the individual protein analysis, pathway analyses between sexes revealed significant enrichment of immune-related processes.

Male mice displayed more abundant alterations over time when compared to female counterparts. Whereas only one pathway was significantly enriched between young and middle-aged, eight pathways were significantly enriched between young and aged timepoints (**Figure 2C and Figure S13, S14**). Of these, *PI3K/Akt signaling* was the only pathway that overlapped between the young vs. middle-aged and the young vs. aged comparisons (**Figure 2D**). This finding suggests *PI3K/Akt signaling* may be a key pathway implicated in aging-induced cartilage degeneration in males, consistent with previous reports demonstrating that *PI3k/Akt signaling* is disrupted in human OA(*16*).

The difference in total normalized perturbation in *PI3K/Akt signaling* from young vs. middle-aged to young vs. aged was only 0.127 (*ΔP_norm_ = P_norm,YoungAged_ – P_norm,YoungMiddle-Aged_*) (**Figure 2D**). Total normalized perturbation is a measure of how much a pathway deviates from physiologic conditions(*17*), with zero being normal physiologic conditions. For these analyses, young mice served as our standard of physiologic conditions (*P_young_ = 0*). This finding suggests that most of the deviation from physiologic conditions is observed from young to middle-aged (**Figure 2B**). *PI3K/Akt signaling* also represented a point of overlap with other pathways that were significantly enriched according to age. For example, *HPV Infection* is predominantly driven by *PI3K/Akt signaling*, *Focal Adhesion,* and *Jak/STAT signaling. PI3K/Akt signaling* is of particular interest given it has been implicated in many chondrocyte cellular processes, including metabolism, apoptosis, and inflammation. Cellular senescence, which was another pathway observed in our analysis, is also a downstream target of *PI3K/Akt signaling* (**Figure 2C)**(*16*).

Further examination of individual proteins associated with *PI3K/Akt signaling* enrichment demonstrated distinctly upregulated or downregulated clusters (**Figure 2E**). GO enrichment analysis revealed that upregulated proteins were associated with *Metabolic Stress*, *Mechano-transduction*, and *Apoptosis*, whereas downregulated proteins were associated with *Matrix Organization*, *Cell Proliferation*, and *Protein Metabolism*. Given the fundamental processes revealed by this GO term analysis, these data suggest that *PI3K/Akt signaling* may be a central regulator of age- and sex-dependent cartilage degeneration (**Figure 2F**). We therefore sought to identify potential age-dependent drivers of this pathway in male mice.

### Age-related decline in α-Klotho is associated with age-induced cartilage degeneration in male mice

Given the predominance of *PI3K-Akt signaling* dysregulation over time in male mice, we probed for candidate proteins that may mediate these changes. There are many regulators of *PI3K-Akt signaling*(*16*). Among these, Insulin signaling is of interest given it has been associated with cellular aging and human chondrocyte health(*18, 19*). Therefore, we probed for potential upstream regulators of the *Insulin-PI3K-Akt* axis. One potential candidate is the longevity protein, α-Klotho (**Figure 2F**). α-Klotho attenuates the onset of tissue aging through a variety of mechanisms related to *PI3K-Akt signaling*, including inhibition of senescence and enhanced autophagy(*20, 21*). While α-Klotho overexpression delays cartilage degeneration in PTOA model(*22*), the role of α-Klotho in age-related KOA has not been clarified. In both mouse and human cartilage, α-Klotho was significantly decreased with increasing age and was significantly associated with more severe cartilage degeneration in male mice (**Figure 3A-C**). In order to evaluate whether α-Klotho plays a direct effect on cartilage health, we evaluated cartilage integrity in mice heterozygously deficient for α-Klotho (Klotho HET). Young and middle-aged Klotho HET male mice displayed significantly accelerated cartilage degradation and increased surface roughness (**Figure 3D, E**). However, such changes were not observed in female young and middle-aged Klotho HET mice (**Figure 3D, E**). These findings suggest that loss of α-Klotho may be a driver of age-related KOA in a sex-dependent manner. Given the limited KOA phenotype in aged female mice, only male mice were used in subsequent experiments.

**Figure 3.**
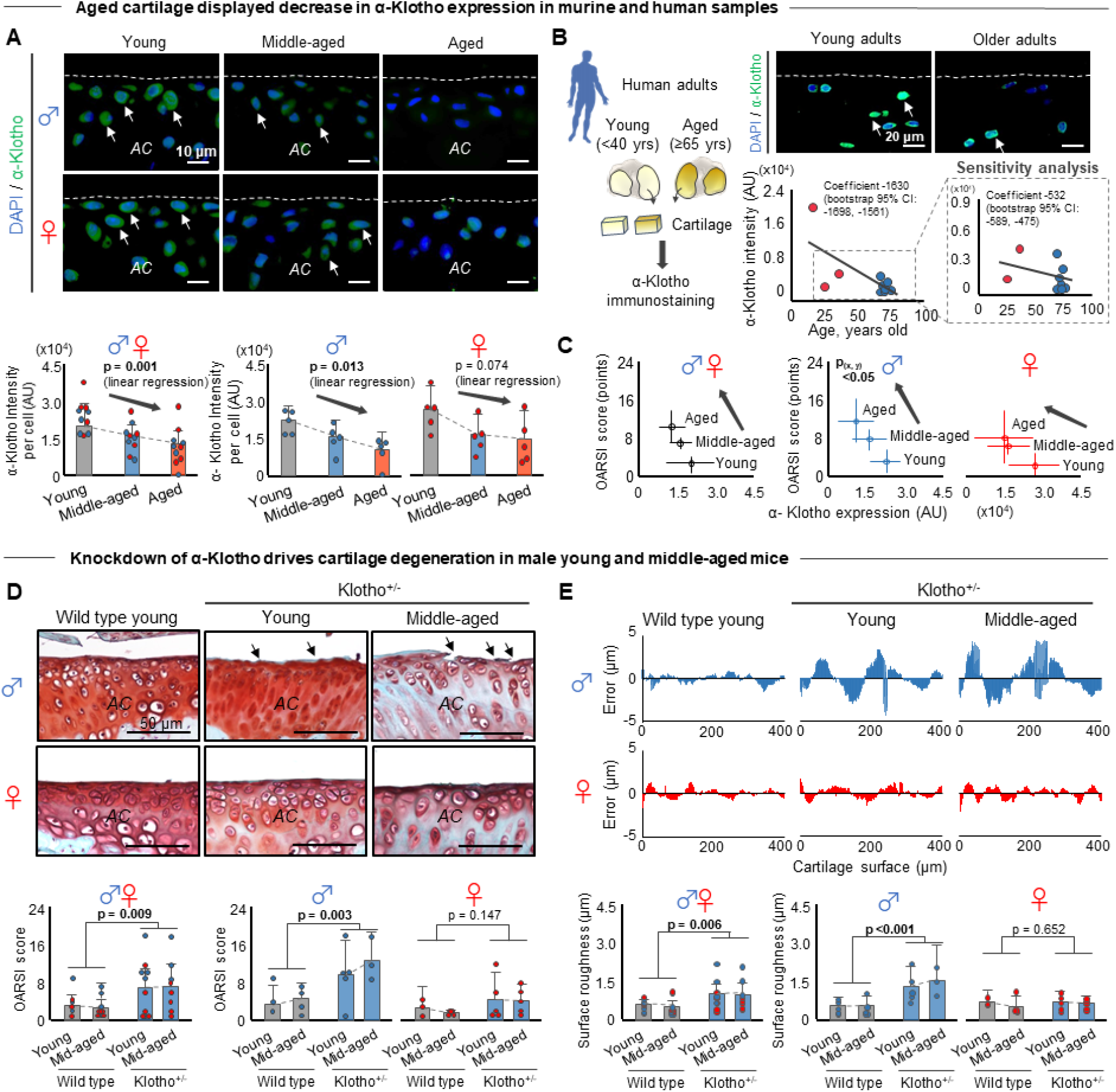
Aging induced progressive α-Klotho decline in cartilage that triggers cartilage degeneration. **A.** Aging induces progressive α-Klotho decline in murine medial tibial plateaus. White arrows indicate α-Klotho-positive chondrocyte. White dash lines indicate cartilage surface. *AC*: articular cartilage. **B.** Cartilage in older adults (≥65 years) displays lower α-Klotho expression than that in young adults (<40 years). **C.** α-Klotho expression decreases with increased cartilage degeneration assessed by OARSI score in a sex-dependent manner. **D.** Loss of function in α-Klotho (Klotho^+/−^) triggers cartilage degeneration in murine medial tibial plateaus in a sex-dependent manner. Black arrows indicate cartilage surface disruption. *AC*: articular cartilage. **E.** Cartilage in Klotho+/−male mice displays higher surface roughness. Statistical analysis was performed using linear regression analysis (**A**, **B**), bootstrap linear regression analysis (**C**), two-way ANOVA (**D**, **E**). P(x,y) <0.05 in Figure B indicates that statistically significant relationship between both aging and α-Klotho decline (x-axis) and aging and OARSI score (y-axis). Data are presented as means ± 95% confidence intervals

### Age-related decline in α-Klotho is associated with alterations in nuclear morphology in mice and humans

*What drives decline in α-Klotho protein levels over time?* Whereas numerous studies in rodents and humans have demonstrated that loss of α-Klotho promotes an aged phenotype in tissues throughout the organism, little is known about the molecular mechanisms driving this decline. It is well known that gene expression can be altered as a result of age-related changes in the cellular microenvironment through mechanotransductive signaling(*23*). Cells sense and respond to signals emanating from the biophysical niche through cytoskeleton-mediated alterations in nuclear morphology(*24*). Indeed, nuclear envelope dysfunction drives chromatin remodeling and has been tightly linked to cellular aging(*25*). The nuclear envelope is primarily composed of nuclear lamina (i.e., lamin A/C and lamin B) and a double membrane through the linker of the nucleoskeleton and cytoskeleton (LINC) complex such as Sun proteins. Lamin A/C is a well-known regulator of nuclear integrity and mechanotransduction(*26, 27*). With this in mind, we revisited the mass spectrometry data to probe for age-related changes in nuclear envelope elements (**Figure 4A**). There was increased expression of nuclear envelope elements including lamin A/C and lamin B2 in aged mice compared to young counterparts (**Figure 4B**). These findings suggest reorganization of the nuclear envelope and resulting pathogenic mechanotransductive signaling may contribute to chondrocyte aging.

**Figure 4.**
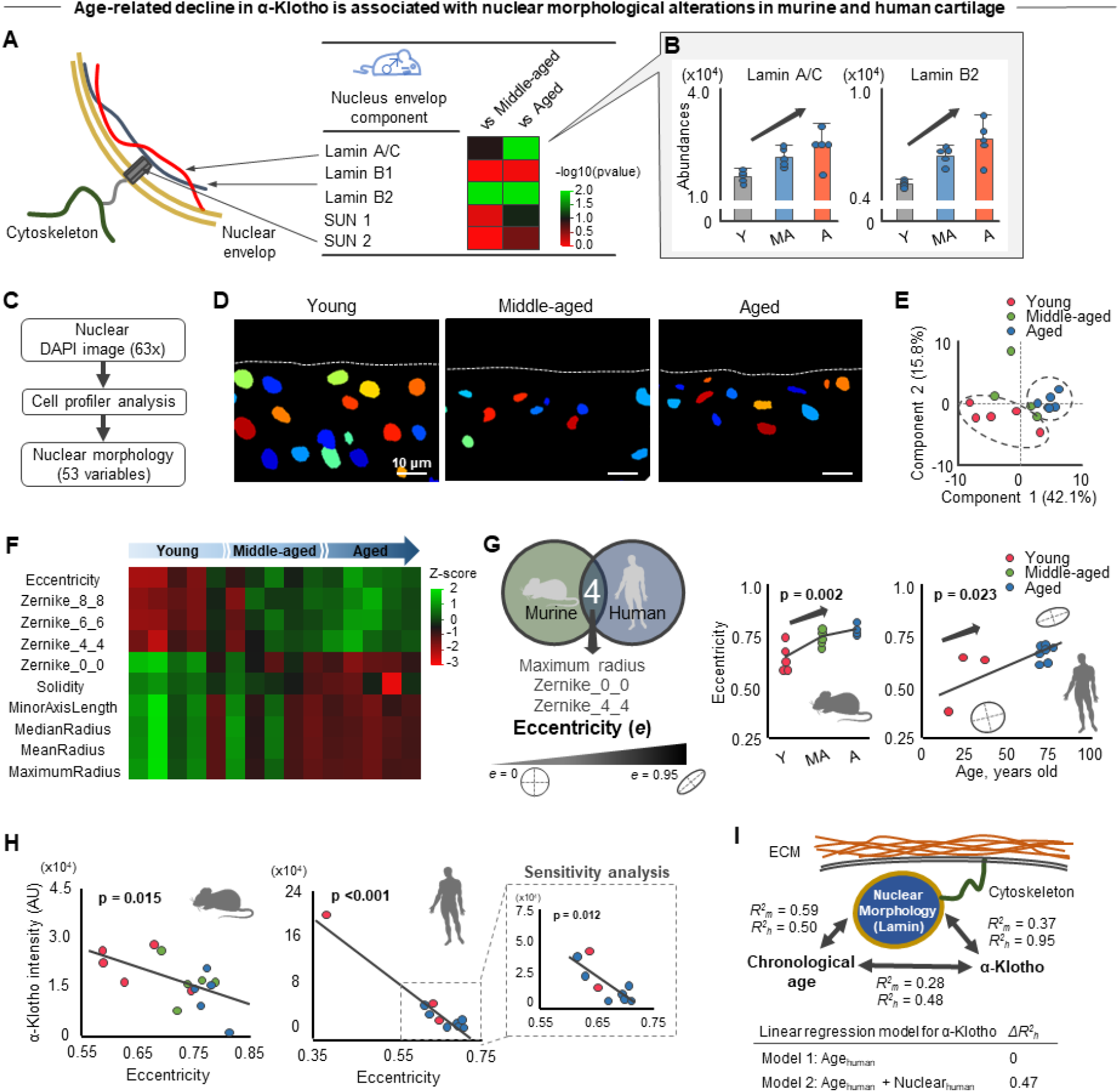
Age-related α-Klotho decline accompanied with nuclear morphological alterations. **A.**Age-related change in proteins associated with nucleus envelop component in male mice. **B.** Aging increases Lamin A/C and B2 protein expression. **C.** Analytical flow of the nuclear morphological analysis (53 variables) using the Cell profiler. **D.** Representative nuclear images in murine medial tibial plateaus generated by Cell profiler. White dash lines indicate cartilage surface. **E.** Principal component analysis (PCA) showing the separate clusters in young and aged nuclear morphology. **F.** Heat map of the top 10 nuclear morphological variables contributing principal component 1, showing the age-related nuclear morphological alterations. Color indicates z-score in each variable. **G.** Murine and human cartilage samples share four nuclear features, in which higher age increases nuclear eccentricity (i.e., less roundness) in murine and human cartilage. **H.** Age-related increased nuclear eccentricity is associated with decreased α-Klotho expression. Sensitivity analysis excluding one outlier data showed a similar trend. **I.** Schematic showing the relationship among chronological age, α-Klotho expression, and nuclear morphology. Linear regression model is provided showing the substantial and indendent contribution of nuclear morphology (Nuclear_human_) in the prediction of α-Klotho expression level in human cartilage beyond the chronological age effect (Age_human_). Coefficient of determinations are provided for mice (*R^2^_m_*) and human cartilage (*R^2^_h_*). Statistical analysis was performed using linear regression analysis (**G, H, I**). Data are presented as means ± 95% confidence intervals.

As a first step to probe this hypothesis, we quantified chondrocyte nuclear morphology in cartilage tissue according to 53 different morphological variables using Cell Profiler software(*28*) (**Figure 4C, D**). Principal component analysis (PCA) of nuclear morphological features revealed clear segregation of nuclei from young and aged chondrocytes **(Figure 4E**). Of the top 10 variables contributing to the first principal component (**Figure 4F**), we identified four common characteristics that changed with aging across both murine and human studies, including maximum radius, Zernike_0_0, Zernike_4_4, eccentricity (**Figure 4G**). Notably, increased nuclear eccentricity (i.e., less spherical) was significantly associated with decreased α-Klotho expression in both murine and human samples (**Figure 4H**). These results suggest that decline in α-Klotho may be attributed, at least in part, to altered nuclear morphology (**Figure 4I**).

### Substrate stiffness regulates a-Klotho expression in chondrocytes regardless of age

Nuclear morphology is mechanically coupled to the surrounding microenvironment where cytoskeletal elements regulate nuclear shape according to matrix stiffness(*29, 30*). As cartilage stiffness in aged mice and humans is 2-3 times higher than young counterparts(*31*), we posited that increased ECM stiffness may drive young chondrocytes towards an aged phenotype. To test this hypothesis, we seeded young or aged primary mouse chondrocytes onto polyacrylamide gels of different stiffness (5kPa, 21kPa, and 100kPa). This stiffness range was selected to mimic a physiologically-relevant ECM stiffnesses in young (5-30 kPa) and aged (50-100 kPa) murine and human knee cartilage(*31*) (**Figure 5A**). This range of stiffness induces significant morphological changes in chondrocytes (**Figure S15**).

**Figure 5.**
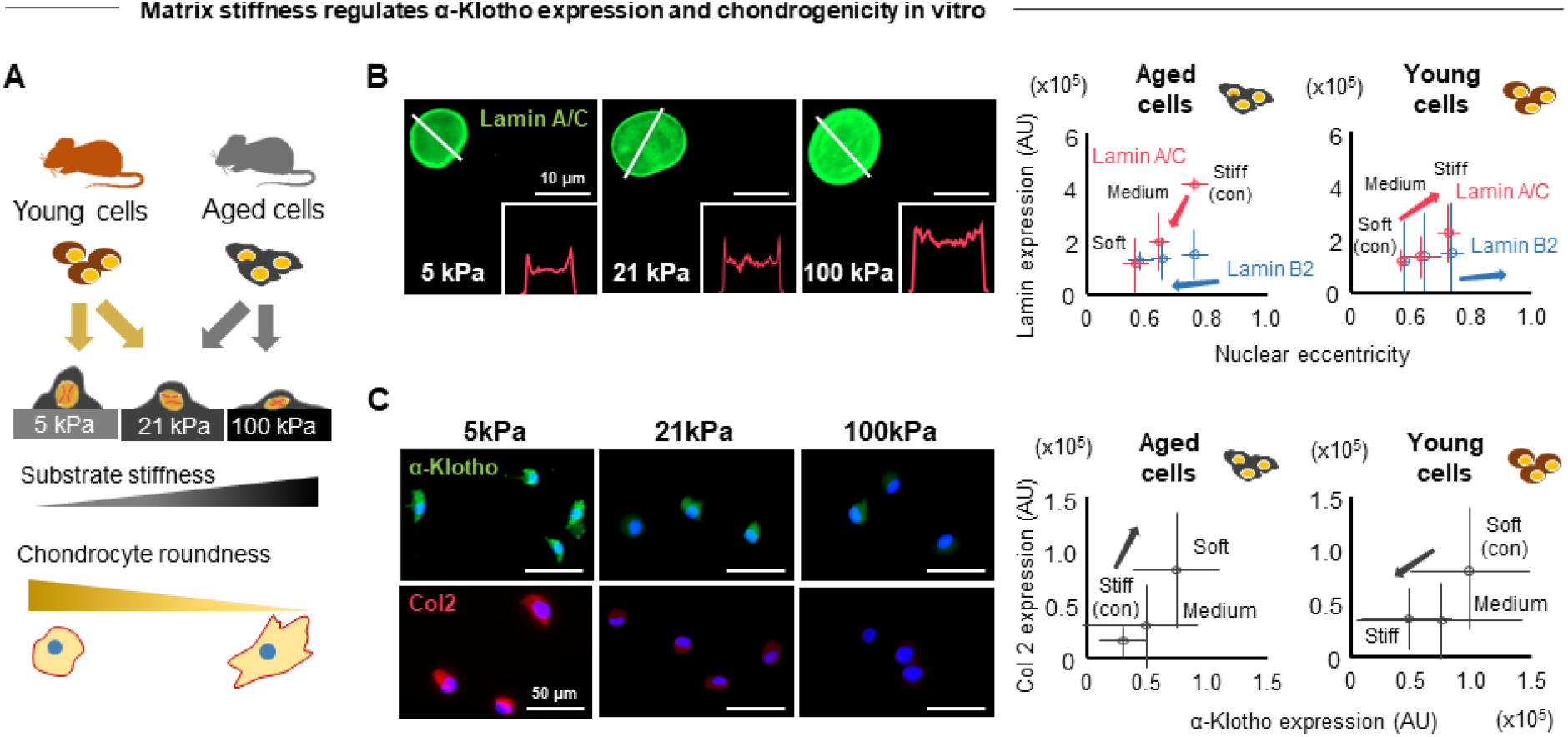
Matrix stiffness regulates chondrocyte α-Klotho expression. **A.** Schematic showing the experimental protocol. Primary chondrocytes isolated from young and aged cartilage were seeded onto polyacrylamide gels engineered to mimic a physiological range of knee cartilage ECM stiffnesses (5kPa, 21kPa, and 100kPa). These range of stiffness differentially influence chondrocytes roundness (see **Figure S15**). **B, C.** Stiff substrate increases expression of Lamin A/C (**B**) and reduces α-Klotho and type II collagen (**C**) in chondrocytes regardless of age of the cell donor. Insets in the fluorescence microscope images highlight increased fluorescence signal in entire nucleus of chondrocytes cultured on stiff substrate. Data are presented as means ± 95% confidence intervals.

Aged chondrocytes cultured on soft substrates displayed a more youthful phenotype when compared to cells cultured on stiffer substrates, as evidenced by lower nuclear lamin A/C expression, a more spherical nuclear shape, as well as increased type II collagen and α-Klotho expression (**Figure 5B, C**). On the other hand, young chondrocytes cultured on stiff substrates developed an aged phenotype, as evidenced by increased lamin A/C expression, a less spherical nuclear shape, and reduced type II collagen and α-Klotho expression (**Figure 5C)**. In contrast, lamin B2 was not sensitive to substrate stiffness (**Figure 5B**). These findings are consistent with a previous report showing that lamin A/C, but not B, is mechanosensitive and influenced by substrate stiffness(*29*). These findings illustrate a mechanistic relationship between ECM biophysical properties, nuclear stiffness, α-Klotho expression, and chondrogenicity.

### Modulation of ECM stiffness increases α-Klotho expression and improves cartilage health in aged mice

To identify ECM proteins that may contribute to the increased nuclear stiffness (i.e., lamin A/C) *in vivo*, we again revisited the mass spectrometry data. Pathway analysis revealed enrichment in ECM-related pathways including *Focal Adhesion*, *ECM Receptor Interaction*, *Hippo signaling,* and *Proteoglycans in Cancer* (**Figure 2C**). PCA further revealed that aged cartilage displayed an altered expression profile in 155 ECM-related proteins detected in our mass spectrometry data (**Figure 6A**). Lysyl oxidase (LOX) emerged within the top 20 proteins contributing to principal component 2, which was positively correlated with lamin A/C (**Figure 6B**). This is particularly of interest given that LOX is one of the major enzymes that induces collagen crosslinking and contributes to PTOA pathogenesis through increased ECM stiffness(*32*). The positive correlation between the LOX and Lamin A/C (**Figure 6C**) and led us to the hypothesis that increased nuclear stiffness and subsequent declines in α-Klotho in aged cartilage may be regulated, at least in part, by LOX.

**Figure 6.**
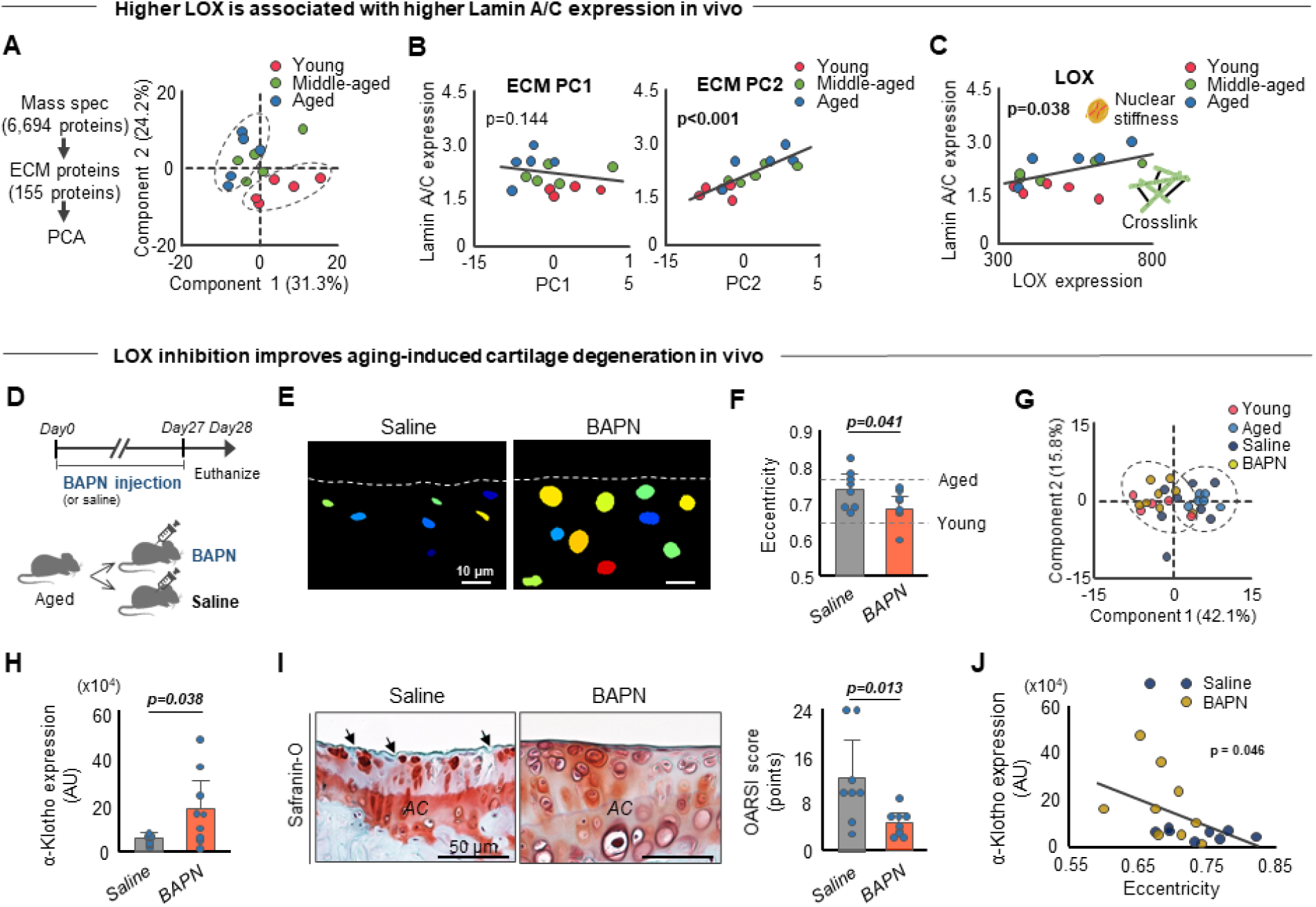
BAPN injection improves α-Klotho expression and cartilage integrity in aged mice. **A.** Principal component analysis (PCA) showing the separate clusters in ECM proteins from young and aged cartilage. **B.** Principal component 2 in ECM protein expression has a link with Lamin A/C expression. **C.** Increased LOX expression is associated with increased Lamin A/C expression, indicating the connection between LOX-mediated increased collagen crosslinking and nuclear stiffness. **D.** Schematic showing the experimental protocol of BAPN daily injection. **E.** Representative Cell Profiler-generated nuclear images in murine medial tibial plateaus after 4-week injection of saline or BAPN. Nuclei were provided with false-colors. White dash lines indicate cartilage surface. **F.** BAPN injection in aged mice decreases nuclear eccentricity towards young level. **G.** PCA showing the same cluster of (1) young and aged+BAPN injection and (2) aged and aged+saline injection. **H.** BAPN injection in aged mice improves α-Klotho expression. **I.** BAPN injection in aged mice improves cartilage integrity. Representative histological sections stained with Safranin-O/Fast Green are provided. Black arrows indicate loss of cartilage matrix. OARSI score (0-24 points; higher value indicates more severe cartilage degeneration) assessed by blinded assessor is provided. **J.** BAPN injection-induced decreased nuclear eccentricity (i.e., more roundness) is associated with higher α-Klotho expression. Statistical analysis was performed using linear regression analysis (**B, C, J**), Student t-test (**G-I**). Data are presented as means ± 95% confidence intervals.

To test this hypothesis and confirm the physiological relevance of *in vitro* findings, we administered β-aminopropionitrile (BAPN), a known inhibitor of LOX, to aged mice daily for four weeks (**Figure 6D**). Histological analysis revealed that cartilage from aged animals treated with BAPN displayed a more spherical nuclear morphology with decreased nuclear eccentricity, thereby mimicking the chondrocyte morphology of young mice (**Figure 6E, F**). PCA analysis further confirmed a more youthful phenotype of chondrocytes from aged mice treated with BAPN (**Figure 6G**). Finally, consistent with our *in vitro* findings, reducing matrix stiffness *in vivo* significantly increased α-Klotho levels in aged chondrocytes and improved cartilage integrity (**Figure 6H- J)**.

## DISCUSSION

The objective of this work was to provide novel and human-relevant insights into pathogenic mechanisms underlying age-related KOA. Through mass spectrometry proteomic analyses, we found that progressive disruption of the intracellular signaling transduction pathway *PI3K/Akt signaling* was associated with the onset of age-related KOA in a sex-dependent manner. In search of upstream candidates that may drive these changes, we discovered that both age-related and genetically-induced declines in α-Klotho, a suppressor of *PI3K/Akt signaling*, drove cartilage degeneration in mice. This age-related decrease in chondrocyte α-Klotho expression was corroborated in young and aged human cartilage. Chondrocyte α-Klotho expression was highly responsive to biophysical characteristics of the ECM. As is observed with increasing age, increased matrix stiffness drove decreased α-Klotho expression. Conversely, reducing matrix stiffness *in vivo* increased α-Klotho expression and improved chondrogenicity of chondrocytes.

### PI3K/Akt signaling: a sex-dependent driver of age-induced cartilage degeneration in mice

Despite the fact that there are many studies characterizing tissue-level changes in cartilage with aging, there is a paucity of work that has thoroughly examined cartilage integrity according to sex. This is a critical shortcoming of the existing literature given that post-menopausal women typically present with more severe KOA(*12*). To address this gap, we thoroughly described the trajectory of cartilage degeneration over time and according to sex using well-established histological analyses(*33*). We found that age-related cartilage degeneration was more severe in male mice, findings that are inconsistent with the clinical evidence of an increased prevalence of KOA in women(*12*). This disconnect may result from the fact that the aged female mice used in our study were non-menopausal, as evidenced by maintained serum estrogen levels across aging. Estrogen affects key signaling molecules in several distinct canonical and non-canonical estrogen signaling pathways, such as PI3K/Akt and PKC/MAPK signaling(*34*). Whereas the beneficial role of systemic estrogen administration in OA development has been investigated *in vivo*, the underlying mechanism remains unknown(*34*). Our data highlight the need for mechanistic studies into the effects of menopause on KOA.

Our findings confirmed previous reports that age-related KOA was accompanied by disruption of the *PI3K/Akt signaling* cascade(*16*). Disruption of *PI3K/Akt signaling* contributes to an imbalance of ECM production and destruction, a pathway that is clearly dysregulated in human OA(*16*). The important role of this pathway in OA development is further evidenced by *in vivo* studies showing that modulating *PI3K/Akt signaling* attenuated pathological cartilage changes in PTOA knees(*35, 36*). Although little is known about the pathogenic contribution of this pathway in age-related KOA, involvement of *PI3K/Akt signaling* is generally in line with previous finding of age-related disruption in insulin-like growth factor 1 signaling in human chondrocytes(*18*). It is noteworthy that alterations in *PI3K/Akt signaling* were already evident in middle-aged mice, coinciding with the onset of cartilage structural abnormalities in humans(*9*). Early disruption of *PI3K-Akt signaling* is supported by a previous microarray analysis showing upregulated inflammation-related genes and down-regulated ECM-related genes in middle-aged mice(*37*), both of which are regulated by *PI3K-Akt signaling*(*16*). The observation that only *PI3K-Akt signaling* was significantly changed in middle-aged mice suggests that this pathway may be an early driver of OA pathogenesis.

### α-Klotho as a potential mediator between alterations in the ECM and cartilage integrity

In search of novel upstream candidates that may regulate *PI3K/Akt signaling*, we found that diminished α-Klotho drives cartilage degeneration. Our finding reinforces previous *in vitro* and *in vivo* studies showing that α-Klotho overexpression counteracts chondrocyte dysfunction and cartilage degeneration(*22, 38*). α-Klotho inhibits insulin growth factor receptor-mediated *PI3K/Akt signaling* and subsequently enhances FoxO, thereby attenuating the deleterious effects of chronic oxidative stress(*21, 39*). FoxO, which increases chondrocyte susceptibility to oxidative stress and impairs cartilage integrity(*40, 41*), is markedly reduced with aging in both murine and human cartilage(*42*). Our results suggest that age-related cartilage degeneration is attributed, at least in part, to reduced protection from oxidative stress by α-Klotho.

Whereas age-related declines in α-Klotho have been linked to the onset of an aged tissue phenotype in many organ systems, our understanding of the mechanisms driving these declines is lacking. The ECM plays a dynamic role in regulating cartilage homeostasis and undergoes extensive remodeling with increasing age, including a decrease in compliance(*31, 43*). It is well established that increased matrix stiffness disrupts chondrocyte functionality via mechanotransductive pathways(*32, 44, 45*), leading to increased cellular senescence. While it is clear that mechanical cues signal alterations in gene expression, the mechanisms by which these cues may drive an aged cellular phenotype are unclear. Here, we identified matrix stiffness as a novel regulator of α-Klotho. Specifically, we found that increased matrix stiffness decreased α-Klotho expression and drove chondrocytes to an aged phenotype *in vitro*. Conversely, decreasing matrix stiffness through inhibition of LOX increased α-Klotho expression and improved cartilage integrity. This finding is in line with previous reports showing that reduced LOX-mediated collagen cross-linking improved cartilage integrity in a PTOA model(*32*). However, this previous study showed LOX-mediated collagen cross-linking to be mediated through mechanotransduction of the RhoA/ROCK axis(*32*) It remains to be seen whether findings from these two studies suggest a previously unappreciated link between RhoA/ROCK signaling and Klotho expression in chondrocytes, or whether the different models may trigger distinct molecular signaling cascades.

One prominent feature linking age-related changes in ECM stiffness to altered α-Klotho expression is nuclear morphological alteration. Remodeling of nuclear morphology is closely associated with modified gene expression and protein synthesis(*46*). In this study, we observed strong relationships between high nuclear deformation and α-Klotho decline in both murine and human samples. Notably, the higher nuclear deformation seen in aged cartilage was recapitulated by a stiff microenvironment. These nuclear morphological alterations are attributed to increased Lamin A/C, a primary mediator of nuclear integrity in response to matrix elasticity compared to Lamin B(*29*). Altered expression of Lamin A/C directly influences chromatin dynamics, leading to altered gene expression and protein synthesis(*47*). Our findings suggest that age-related alterations in ECM biophysical properties change nuclear integrity, ultimately inhibiting α-Klotho expression. Studies have shown that looser packed (i.e. more transcriptionally active) chromatin is more subject to changes in cellular forces than more condensed (i.e. less transcriptionally active) chromatin(*48*). Through these investigations, the mechanotransductive pathway may be expanded to include regulation of α-Klotho expression and ultimate inoculation of an aged phenotype, which is a necessary direction of future studies.

Taken together, the studies presented suggest a novel molecular mechanism for age-related KOA. We found biophysical alterations in cartilage ECM disrupt chondrocyte nuclear stiffness and decrease α-Klotho expression. In turn, we found decreased α-Klotho compromised chondrocyte homeostasis and tissue integrity, ultimately driving OA. While future studies are needed to fully appreciate the capacity of the ECM-α-Klotho-PI3K/Akt axis, the coalescion of our findings provides a springboard for a new generation of therapeutics targeting the insidious nature of age-related KOA.

## METHODS

### Animals

Experiments were performed using young (4-6 months; body mass), middle-aged (10-14 months; body mass), and aged (21-24 months; body mass) male and female C57/BL6 mice, as well as young (4-6 months) and middle-aged (10-14 months) male and female Klotho heterozygotes mice (Kl^+/−^; B6; 129S5-Kltm1-Lex, 7-10 months, UC Davis). Mice were obtained from the NIA Rodent Colony and Jackson Laboratories. Prior to inclusion in experiments, animals were evaluated and those with visible health abnormalities were excluded. Mice were housed in cages holding an average of 3-4 mice per cage with a temperature-controlled environment and 12-h light/dark cycles. The animals can access food and water ad libitum. All animal experiments were performed in accordance with the ARRIVE guidelines(*49*) and approved by the University of Pittsburgh’s Institutional Animal Care and Use Committee.

### Human cartilage samples

Normal human articular cartilage tissue in knee joint from young (<40 years old; n = 3) and older (≥65 years old; n = 7) donors were obtained through the National Disease Research Interchange (Philadelphia, PA) with approval from the University of Pittsburgh Committee for Oversight of Research and Clinical Training Involving Decedents (CORID). Cartilage samples were fixed by 10% buffered formalin phosphate (Fisher Chemical, Fair Lawn, NJ, USA) overnight and dehydrated with 50%, 70%, 95% and 100% ethanol on the second day. After being treated in xylene for 2 hours and then in liquid paraffin overnight, samples were embedded in paraffin. The sample blocks were sectioned at a thickness of 6 μm thickness using a Leica microtome (Model RM 2255). For DAPI staining, sections were deparaffinized using Histo-Clear II (National Diagnostics, Atlanta, GA, USA), rehydrated, and stained with DAPI mounting medium (Antifade Mounting Medium with DAPI H-1200-10, Vector Laboratories, Burlingame, CA, USA)

### BAPN injection of mice

Aged male C57/BL6 mice (21-24 months; body mass) received daily subcutaneous injections of either saline, as a control, or BAPN (290 μg/μl in saline) for 4 weeks every day. Volume of the injection (40-80 μl) was determined based on mice body mass (*m*) with following protocol: 40ul (18.45g≤*m*<23.73g), 50μl (23.73g≤*m*<29.00g), 60μl (29.00g≤*m*<34.27g), 70μl (34.27g≤*m*<39.55g), 80μl (39.55g≤*m*<44.82g). Mice were weighed every other day beginning on day 0 and ending on day 28 to ensure proper dosage of BAPN.

### Serum collection

We collected serum from animals under anesthesia by isoflurane in accordance with established protocol(*50*). After collection, the animals were euthanized via cervical dislocation. The collected blood was allowed to clot in a 2 mL tube at room temperature for one hour. Then, the blood was centrifuged for 30 minutes at 13,000 rpm at 4 °C in a microcentrifuge. The serum was collected and aliquoted into 50 μl tubes and stored at −20 °C until used. Any samples displaying hemolysis (as evidenced by pink/red coloration) were not included in the analysis.

### Estrogen ELISA

The Estrogen ELISA was conducted according to protocol using the 17 beta Estradiol ELISA Kit (ab108667, Abcam). This kit was validated by Marino FE *et al.*(*51*). Briefly, 25 μl of blood serum samples, prepared standards, or controls were added in duplicates to a 96 well plate. 200 μl of the 17 beta Estradiol-HRP Conjugate were added to each well. This plate was then incubated at 37°C for two hours. Samples, standards, and controls were aspirated, and wells were washed three times with 300 μl of diluted washing solution (soak time > 5 seconds) using an automated plate washer (BioTek 50TS). After washing, 100 μl tetramethylbenzidine (TMB) Substrate Solution was added to each well, and the plate was incubated for 30 minutes in the dark at room temperature. After incubation, 100 μl of Stop Solution was added into all wells in the same order and at the same rate as the substrate solution. Absorbance was measured at 450 nm with Spectramax M3 plate reader (Molecular Devices) within 30 minutes of adding the Stop Solution. Construction of standard curve and subsequent analyses were performed in Microsoft Excel.

### Histological preparation and semi-quantitative histological score for cartilage degeneration

Mice knee joints and human cartilage samples were fixed in 4% paraformaldehyde overnight at 4°C, and decalcified in 20% ethylene diamine tetra acetic acid solution for 10 days. Decalcified paraffin sections (5 μm thickness) were prepared from central region of the mice knee joints in the frontal plane in accordance with the OARSI recommendation(*52*). Decalcified paraffin sections from human cartilage specimen were also prepared. The paraffin sections were stained with Safranin-O/Fastgreen/Hematoxilin to evaluate the severity of cartilage lesions. This study focused on medial tibial cartilage given this is the region most typically affected in humans(*8*) and its severity is typically equal to or higher than those for the femoral condyle in the majority of aged mice(*53*). The OARSI scoring system, consisting of six grades and four stages on a scale from 0 (normal) to 24 (severe cartilage lesion), was used for semi-quantitative evaluation of cartilage lesion severity(*33*). The most severe score was evaluated as the maximum OARSI score. We did not consider adding the summed scores generated from different sections, as discriminative ability for age-related cartilage degeneration is comparable to the maximum score(*53*). A trained examiner (AB) performed grading in each histological section in a blinded manner.

### Chondrocytes isolation

Primary mice chondrocytes were isolated in accordance with an established protocol(*54*). Briefly, cartilage tissue was harvested from femoral head, femoral condyle, and tibia cartilage using small scissors and tweezers while ensuring that minimal fat, muscle, ligament, and tendons were included in the tissue harvest. After brief wash by PBS two times, cartilage pieces were digested with 0.1% (w/w) type II collagenase (cat no. 4176, Worthington Biochemical corp., NJ) in low-glucose DMEM (cat no. 11965-092, Gibco) with 10% (vol/vol) FBS (cat no. SH30070.03, Hyclone) and 1% (vol/vol) Pen/Strep at 37℃ for overnight under 5%CO_2_ in a petri dish. After the digestion, the filtrate was passed through a 50-μm strainer and cells were culture with growth media containing DMEM supplemented with 10% FBS and 1% Pen/Strep until the cell reached to confluent. First-passage cells were used for all the experiments. The isolated cells were characterized by type II collagen immunofluorescence, and over 95% cells were type II collagen positive.

### Preparation of fibronectin-coated pAAm substrates

We prepared pAAm gels with different stiffness (5 kPa, 21 kPa and 100 kPa) in accordance with a previous study(*55*). The pAAm gels were made on glass coverslips, which are pre-treated with 0.1 N sodium hydroxide (cat. SS255-1, Fisher Scientific, IL), 0.5% 3-aminopropyltrimethoxysilane (cat no. AC313251000, Acros Organics, Belgium) and 0.5% glutaraldehyde (cat no. BP25481, Fisher Scientific, IL) to improve the gel adhesion. The detailed gel compositions were provided in **Table S1**. To facilitate the cell adhesion, the surfaces of prepared hydrogels were further conjugated with fibronectin (100 μg/mL, from bovine plasma, Sigma) by using sulfo-SANPAH (cat no. NC1314883, Proteochem Inc., UT) as the crosslinker. Prior to seeding cells, gels were UV-sterilized in a cell culture hood for 30 minutes. Gels were kept hydrated in HEPES or PBS during all preparation steps.

### Cell fate influenced by matrix stiffness

Isolated primary chondrocytes from young and aged mice were plated (5000 cells per mm^2^) on 5kPa, 21kPa, and 100kPa pAAM gel and cultured in low-glucose DMEM supplemented with 10%FBS and 1% Pen/Strep at 37℃ under 5%CO_2_. On day 3, the culture medium was removed and fixed by 4% PFA for 10 minutes. After a triple wash by PBS, cells were kept in PBS at 4°C until used.

### Immunofluorescence and imaging

Immunofluorescence analysis for tissue section and/or cell-seeded pAAm gel was performed to determine the signal intensity of type II collagen, α-Klotho, Lamin A/C, and Lamin B2 in accordance with established protocol(*50*). Briefly, after a triple wash by PBS, cells were permeabilized with 0.1% triton-X (Fluka 93420) for 15 minutes, followed by a one-hour blocking step using 0.1% triton-X with 3% Bovine Serum Albumin (BSA, Sigma A7906) in PBS. The cells were then incubated with primary antibodies overnight at 4°C. A similar process was followed for the decalcified tissue paraffin. For antigen-retrieval for the decalcified tissue paraffin section, they were incubated in sodium citrate buffer for 2 hours at 60°C before the blocking step. After the blocking step, the tissue or cells were incubated overnight at 4°C with the following primary antibodies in antibody solution (0.1% Triton-X+3% BSA+5% Goat Serum), at the dilutions provided in **Table S2**. One negative control slide per staining set was generated by deleting the primary antibody in the antibody solution.

After a triple wash by PBS, the samples were incubated with host-specific secondary antibodies conjugated with Alexa Fluor 488 (Fisher Scientific) in antibody solution for one hour at room temperature at the dilutions of 1:500. Following a triple wash with PBS, the samples were stained with DAPI for 2 minutes and then washed with PBS again. The samples were mounted with coverslips using Gelvatol mounting medium (Source: Center of Biologic Imaging, University of Pittsburgh). The antibody of α-Klotho antibody (R&D systems, MAB1819, Lot# KGN0315101) was validated for skeletal muscle histological section in our previous study(*50*). We also validated the same α-Klotho antibody using the decalcified paraffin section from a wild type and a Kl−/− mouse knee joint. We observed minimal background staining in the Kl−/− section as compared to the wild-type counterparts.

Slides were imaged using a Zeiss Observer Z1 semi-confocal microscope. All images were collected at 20x or 63x magnification. Negative control slides were used to the threshold for the signal intensity and to set the exposure time for individual channels. All images for quantitative analysis in a given experiment were taken under the same imaging conditions. Fluorescence intensity was quantified using Image J

### Quantification of cellular and nuclear morphology

DAPI and F-actin images were obtained at 63x and 20x magnification using a Zeiss Observer Z1 semi-confocal microscope, respectively. Afterward, image processing and morphome feature extraction were performed using CellProfiler software (v4.0, The Broad Institute)(*28*). Fifty-three shape features of cells and nuclei were determined using the “identify primary objects” followed by the “measure object size shape” and “export to spreadsheet” module. Principal Component Analysis (PCA) was performed for the data reduction identifying the principal components (PCs) that represent the differences in the cellular and nuclear morphology. To determine variables of cellular and nuclear shape contributing to PCs, loading matrix, a correlation between the original variables and PCs was extracted.

### LC/MS-MS mass spectrometry-based proteomics

Knee cartilage from male and female young, middle-aged, and aged C57/BL6 mice were microdissected as detailed by previous study(*56*). Briefly, cartilage tissue was harvested from femoral condyle and tibial plateau using small scissors and tweezers while ensuring that minimal fat, muscle, ligament, and tendons were included in the tissue harvest. Immediately after dissection, cartilage samples were washed with PBS. Samples were then lyophilized overnight and stored at-80℃ until shipment to the Proteome Exploratory Laboratory at Cal Tech.

Cartilage samples from each knee were lysed in 150 μl 8M urea/100mM TEAB by grinding for 1 min with 0.5 mL size tissue grinder pestles (Fisher Scientific #12141363), tip sonication with a Fisher Scientific 550 Sonic Dismembrator on ice at 20% power using cycles of 20 sec on/20 sec off for 4 minutes total, followed by another grinding step for 1 min. Samples were then clarified by centrifugation at 16,000g for 5 minutes at room temperature and the lysate collected for protein quantitation by BCA assay (Pierce). Each lysate was then reduced with 1.12 μl 500mM TCEP for 20 min at 37°C and alkylated with 3.36 μl 500mM 2-Chloroacetamide for 15 min at 37°C in the dark. Samples were then digested with a 1:200 ratio of LysC to lysate for 4 hr at 37°C, followed by dilution with 450 μl 100mM TEAB, addition of 6 μl 100mM CaCl2, and digestion overnight (16 hr) with 1:30 Trypsin at 37°C. Digestions were stopped by acidifying with 20 μl 20% TFA, desalted on C18 spin columns (Pierce #89870) according to manufacturer instructions, and lyophilized to dryness. Peptides were then resuspended in 50 μl 0.1% formic acid and peptide amounts measured with the Pierce Quantitative Colorimetric Peptide Assay.

15 μg peptides from each sample were lyophilized, resuspended in 50 μl 100mM TEAB, labeled with 0.25 mg TMTpro reagents dissolved in 10 μl anhydrous acetonitrile for 1 hr at room temperature, and quenched with 2.5 μl 5% hydroxylamine for 15 min at room temperature. Concurrently, a single 30 μg pooled bridging control sample was created by mixing 1 ug from each of the 30 samples into a single tube which was then lyophilized, resuspended in 100 μl 100mM TEAB, labeled with 0.5 mg TMTpro-134 dissolved in 20 μl anhydrous acetonitrile for 1 hr at room temperature, and quenched with 5 μl 5% hydroxylamine for 15 min at room temperature. All 15 male samples were then combined into one sample and all 15 female samples were combined into a second sample. Half of the labeled bridging control sample was then mixed into each and the two resulting mixed samples were lyophilized and stored at-20°C. 100 ug of each sample was then fractionated with the Pierce High pH Reversed-Phase Peptide Fractionation Kit (Thermo #84868) according to manufacturer instructions and the resulting 8 fractions were lyophilized. Each fraction was resuspended in 20 μl 0.2% formic acid and peptide quantitation performed with the Pierce Quantitative Colorimetric Peptide Assay. Fractions 7 and 8 from both samples had very low peptide amounts and were thus combined with that sample’s fraction #6 for a total of 6 fractions per sample.

Liquid chromatography-mass spectrometry (LC-MS) analysis of peptide fractions was carried out on an EASY-nLC 1000 coupled to an Orbitrap Eclipse Tribrid mass spectrometer (Thermo Fisher Scientific). For each fraction, 1 μg peptides were loaded onto an Aurora 25cm x 75μm ID, 1.6μm C18 reversed phase column (Ion Opticks, Parkville, Victoria, Australia) and separated over 136 min at a flow rate of 350 nL/min with the following gradient: 2–6% Solvent B (7.5 min), 6-25% B (82.5 min), 25-40% B (30 min), 40-98% B (1 min), and 98% B (15 min). MS1 spectra were acquired in the Orbitrap at 120K resolution with a scan range from 350-1800 m/z, an AGC target of 1e6, and a maximum injection time of 50 ms in Profile mode. Features were filtered for monoisotopic peaks with a charge state of 2-7 and a minimum intensity of 2.5e4, with dynamic exclusion set to exclude features after 1 time for 45 seconds with a 5-ppm mass tolerance. HCD fragmentation was performed with collision energy of 32% after quadrupole isolation of features using an isolation window of 0.5 m/z, an AGC target of 5e4, and a maximum injection time of 86 ms. MS2 scans were then acquired in the Orbitrap at 50K resolution in Centroid mode with the first mass fixed at 110. Cycle time was set at 3 seconds.

Analysis of LCMS proteomic data was performed in Proteome Discoverer 2.5 (Thermo Scientific) utilizing the Sequest HT search algorithm with the mouse proteome (UniProt UP000000589; 55,485 proteins covering 21,989 genes with a BUSCO assessment of 99.8% genetic coverage). Search parameters were as follows: fully tryptic protease rules with 2 allowed missed cleavages, precursor mass tolerance set to 20 ppm, fragment mass tolerance set to 0.05 Da with only b and y ions accounted for. Modifications were considered static for TMTpro (K, N-term), with dynamic modifications considering Loss of Protein N-term Methionine, Acetyl (N-term), Oxidation (M), Carbamidomethyl (C) and Phosphoration (S, T). Percolator was used as the validation method, based on q-value, with a maximum FDR set to 0.05. GO Biological Process terms were generated via Proteome Discoverer Sequest HT algorithm.

Quantitative analysis is based on TMT MS2 reporter ions generated from HCD fragmentation, with an average reporter S/N threshold of 10, used a co-isolation threshold of 50 with SPS mass matches set at 65%. Normalization was performed at the peptide level, and protein ratios were calculated from the grouped ion abundances, with protein FDR set to a maximum of 0.05.

The mass spectrometry proteomics data have been deposited to the ProteomeXchange Consortium via the PRIDE partner repository(*57*) with the dataset identifier PXD024062 and 10.6019/PXD024062 (Username, Password: 2pCZoIkT). Kyoto Encyclopedia of Genes and Genomes pathway analyses were performed using ROnToTools R Code(*58*). We chose this program for its ability to integrate both over-representation analyses and functional class scoring. The code was used as written, with the exception of changing ‘hsa’ to ‘mmu’ such that mouse pathways were referenced. Our dataset matrices were generated in Excel following the format provided in the sample dataset(*58*).

### Statistical analysis

All statistical analyses were performed using JMP Pro 14 software (SAS Institute, Cary, NC). The data are displayed as means, with uncertainty expressed as 95% confidence intervals (mean ± 95% CI). Linear regression analyses or two-way ANOVA were performed. We checked the features of the regression model by comparing the residuals vs. fitted values (i.e., the residuals had to be normally distributed around zero), and independence between observations. No correction was applied for multiple comparison because that these outcomes determined *priori* and outcomes were highly correlated. Any statistical analyses considered confounder (e.g., body mass) because of the small sample size. We conducted a complete-case analysis in the case of missing data. In all experiments, p-values <0.05 were considered statistically significant. Throughout this text, “*n*” represents the number of independent observations of knees or cells from different animals.

### Methodological rigor

This study was conducted according to the ARRIVE essential 10(*59*). Where possible, power analysis from pilot data was done to select the number of animals needed for the study using Power and Sample Size Program (version 3.1.2; Vanderbilt University Medical Center, Nashville, TN)(*60*). For example, sample size calculation estimated that 10 mice were required to get the statistical power of 0.8 based on the histology score from aging cohort (n = 5 in male young and male aged mice). In addition, a priori power analysis for mass spectrometry estimated that five mice in each group give us a statistical power of 0.90 when a two-fold change is detected, assuming 20% variation(*61*). For *in vivo* experiments including BAPN injection, animals were randomly allocated into BAPN and saline (control) groups using a computer-generated randomization. BAPN and saline treatments were conducted in the same condition and the order of treatment was randomly performed. We randomly replaced the cage location to prevent any bias from environment. All histological outcome assessments were conducted in a blinded manner.

## Acknowledgments

This study was supported in part by (1) a Grant-in-Aid from the Japan Society for the Promotion of Science for Overseas Research Fellowships for HI, (2) NIA R01AG052978, NIA R01AG061005, and NIA P2CHD086843 for FA, and (3) the National Institute of General Medical Sciences of the National Institutes of Health under Award Number T32GM008208 for GG. The funders had no role in study design, data collection and analysis, decision to publish, or preparation of the manuscript.

## Data availability

The datasets used and/or analyzed during the current study are available from the corresponding author on reasonable request.

