## Supplementary Appendix for "Age-related increase in matrix stiffness downregulates α-Klotho in chondrocytes and induces cartilage degeneration"

**The PDF file includes:**

Figure S1. Female Aged vs. Young Mass Spectrometry Individual Protein Volcano Plot
Figure S2. Female Middle-Aged vs. Young Mass Spectrometry Individual Protein Volcano Plot
Figure S3. Male Aged vs. Female Aged Mass Spectrometry Individual Protein Volcano Plot
Figure S4. Male Middle-Aged vs. Female Middle-Aged Mass Spectrometry Individual Protein Volcano Plot
Figure S5. Male Young vs. Female Young Mass Spectrometry Individual Protein Volcano Plot
Figure S6. Male Aged vs. Young Mass Spectrometry Individual Protein Volcano Plot
Figure S7. Male Middle-Aged vs. Young Mass Spectrometry Individual Protein Volcano Plot
Figure S8. Female Aged vs. Young ROnToTool Pathway Analysis Statistics
Figure S9. Female Middle-Aged vs. Young ROnToTool Pathway Analysis Statistics
Figure S10. Male Aged vs. Female Aged ROnToTool Pathway Analysis Statistics
Figure S11. Male Middle-Aged vs. Female Middle-Aged ROnToTool Pathway Analysis Statistics
Figure S12. Male Young vs. Female Young ROnToTool Pathway Analysis Statistics
Figure S13. Male Aged vs. Young ROnToTool Pathway Analysis Statistics
Figure S14. Male Middle-Aged vs. Young ROnToTool Pathway Analysis Statistics
Figure S15. Chondrocytes morphological changes in different stiffness of polyacrylamide gels

Table S1. Preparation and characterization of pAAm hydrogels
Table S2. Information and dilution protocol of antibodies

This supplementary material has been provided by the authors to give readers additional information about their work.

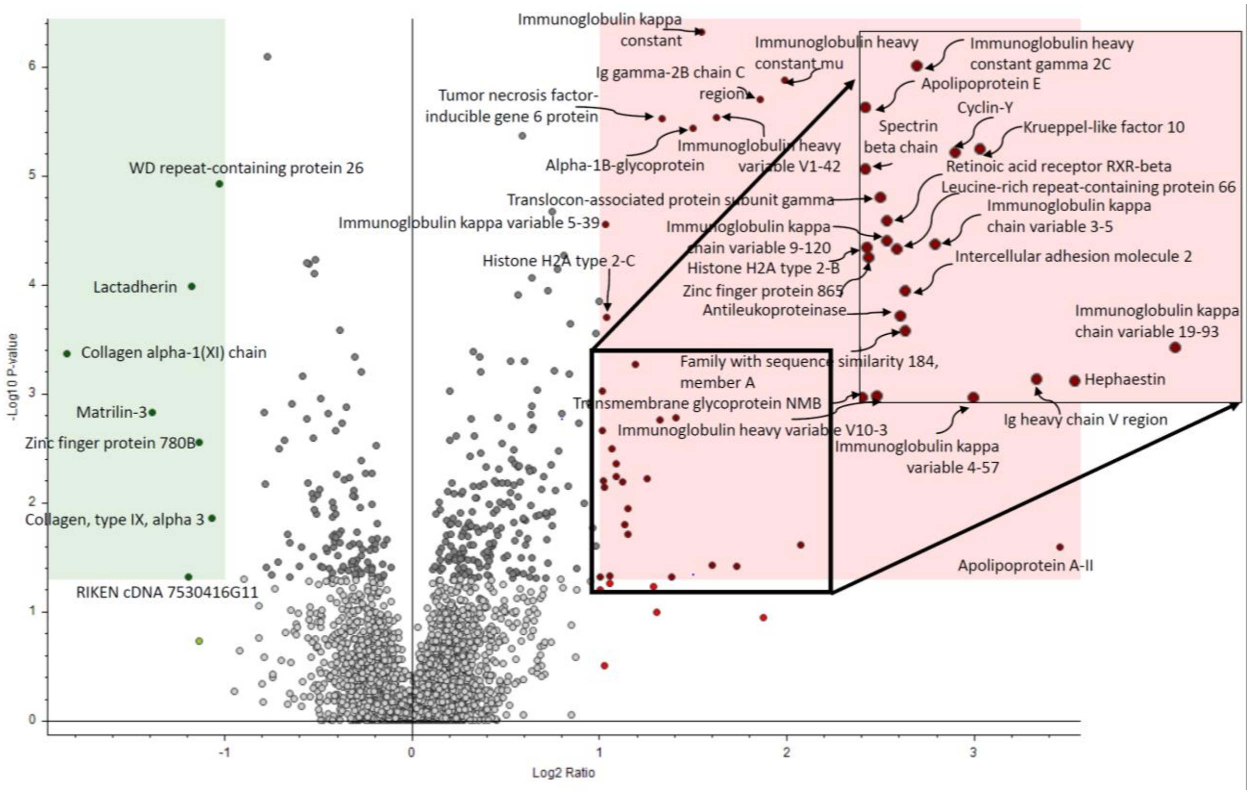


**Figure S1. Female Aged vs. Young Mass Spectrometry Individual Protein Volcano Plot**

Individually identified proteins were determined to be significantly different if the $\log_2$(ratio of female aged vs young) was greater than 1 or less than -1 and the $-\log_{10}$(p-value) was greater than 1.25.

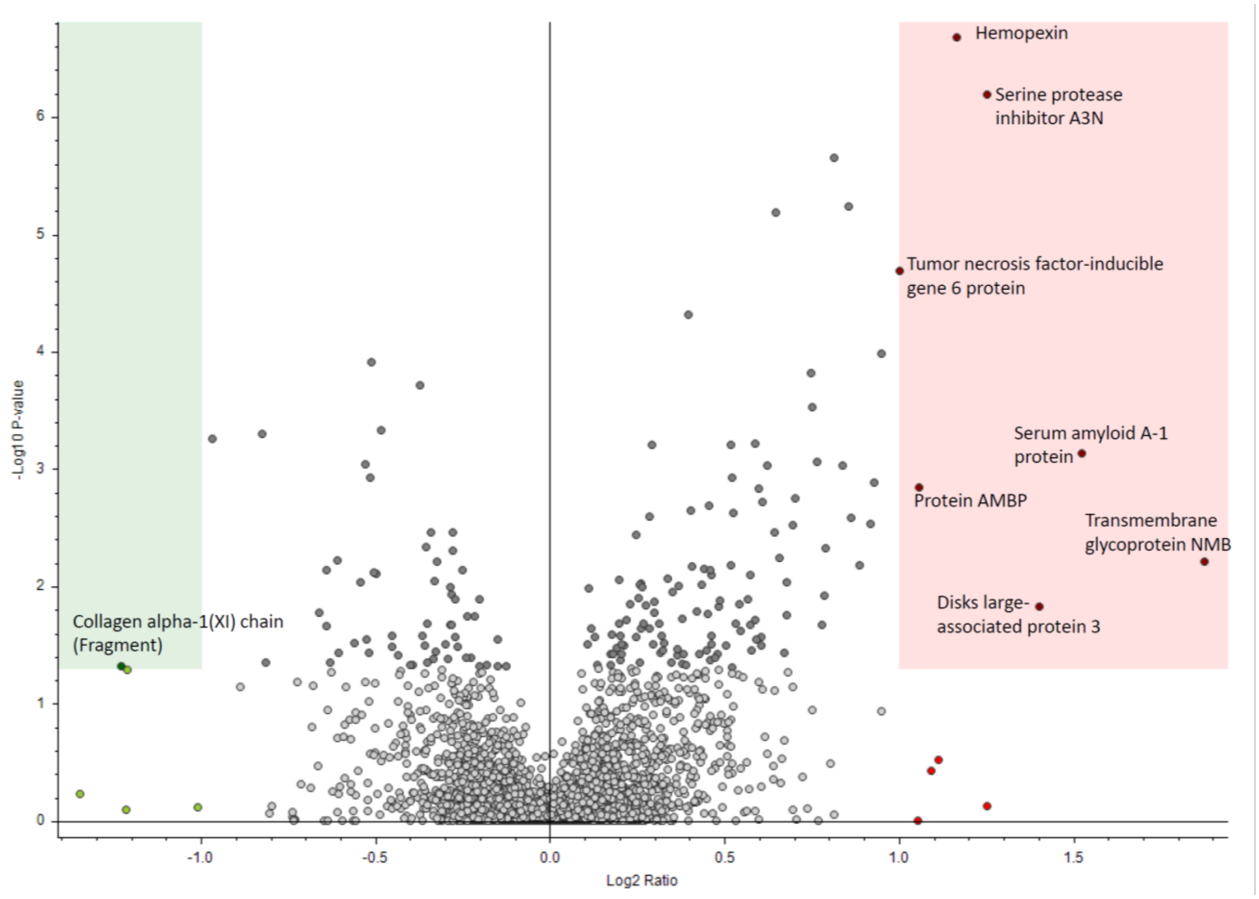


**Figure S2. Female Middle-Aged vs. Young Mass Spectrometry Individual Protein Volcano Plot**
Individually identified proteins were determined to be significantly different if the $\log_2$(ratio of female middle-aged vs young) was greater than 1 or less than -1 and the $-\log_{10}$(p-value) was greater than 1.25.

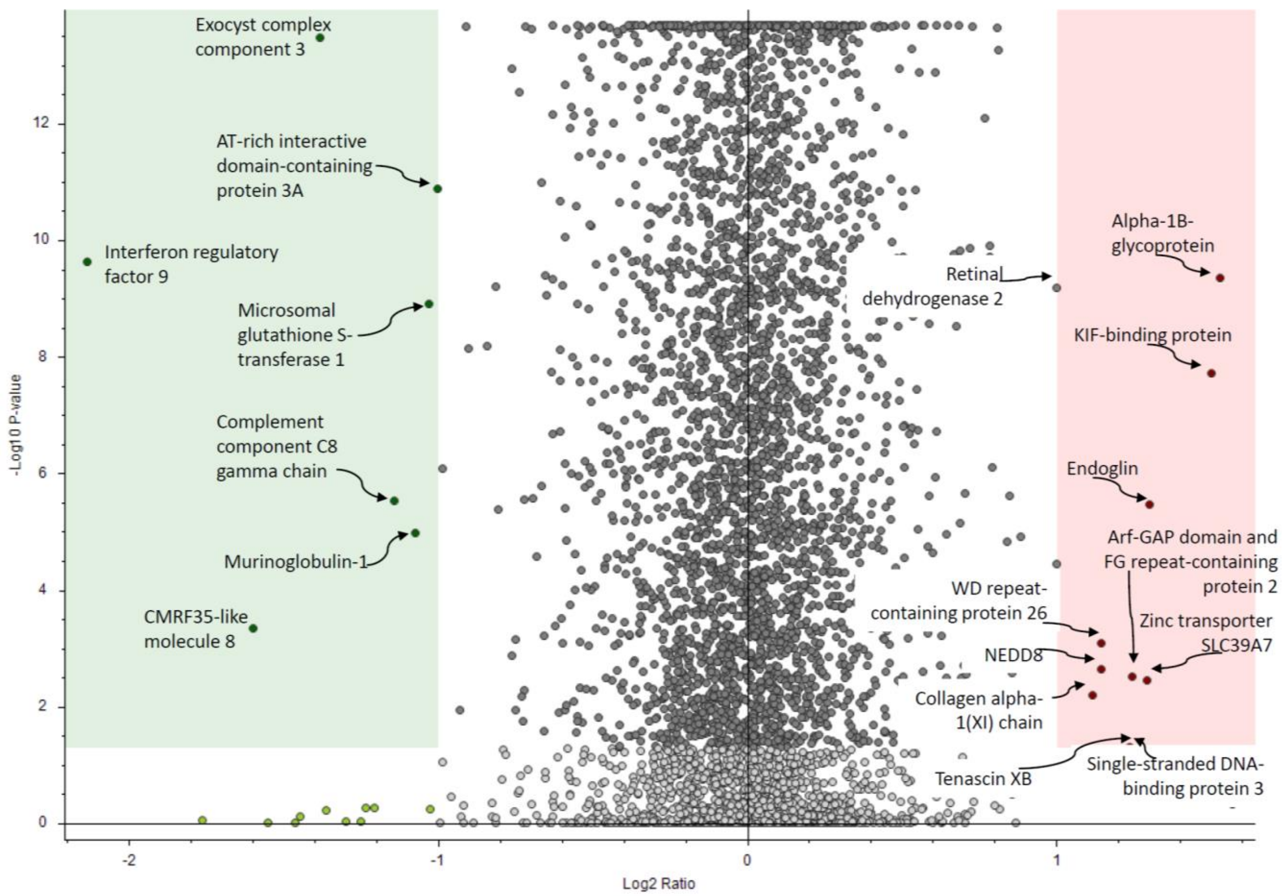


**Figure S3. Male Aged vs. Female Aged Mass Spectrometry Individual Protein Volcano Plot**

Individually identified proteins were determined to be significantly different if the $\log_2$(ratio of male aged vs. female aged) was greater than 1 or less than -1 and the $-\log_{10}$(p-value) was greater than 1.25.

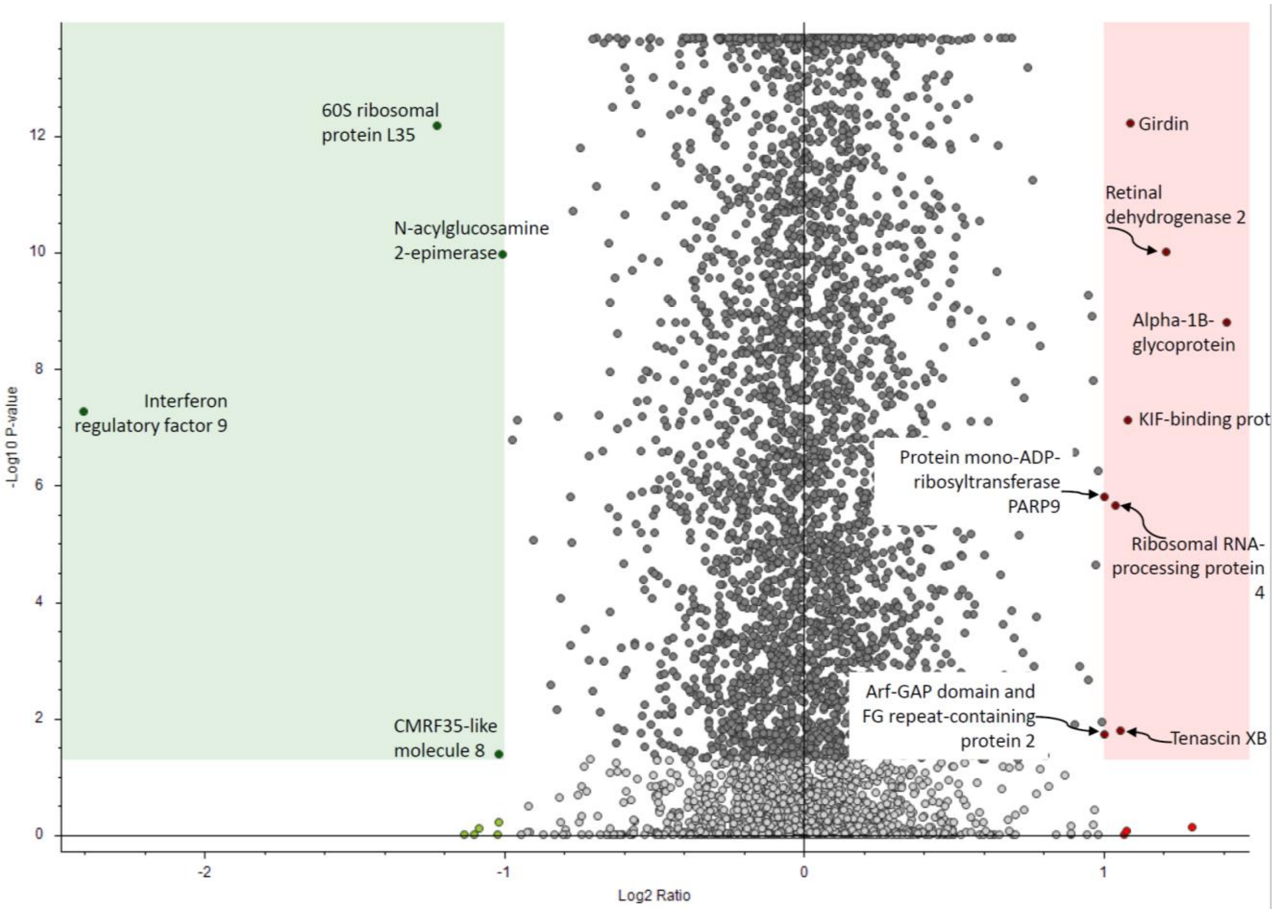


**Figure S4. Male Middle-Aged vs. Female Middle-Aged Mass Spectrometry Individual Protein Volcano Plot**

Individually identified proteins were determined to be significantly different if the $\log_2$(ratio of male middle-aged vs. female middle-aged) was greater than 1 or less than -1 and the $-\log_{10}$(p-value) was greater than 1.25.

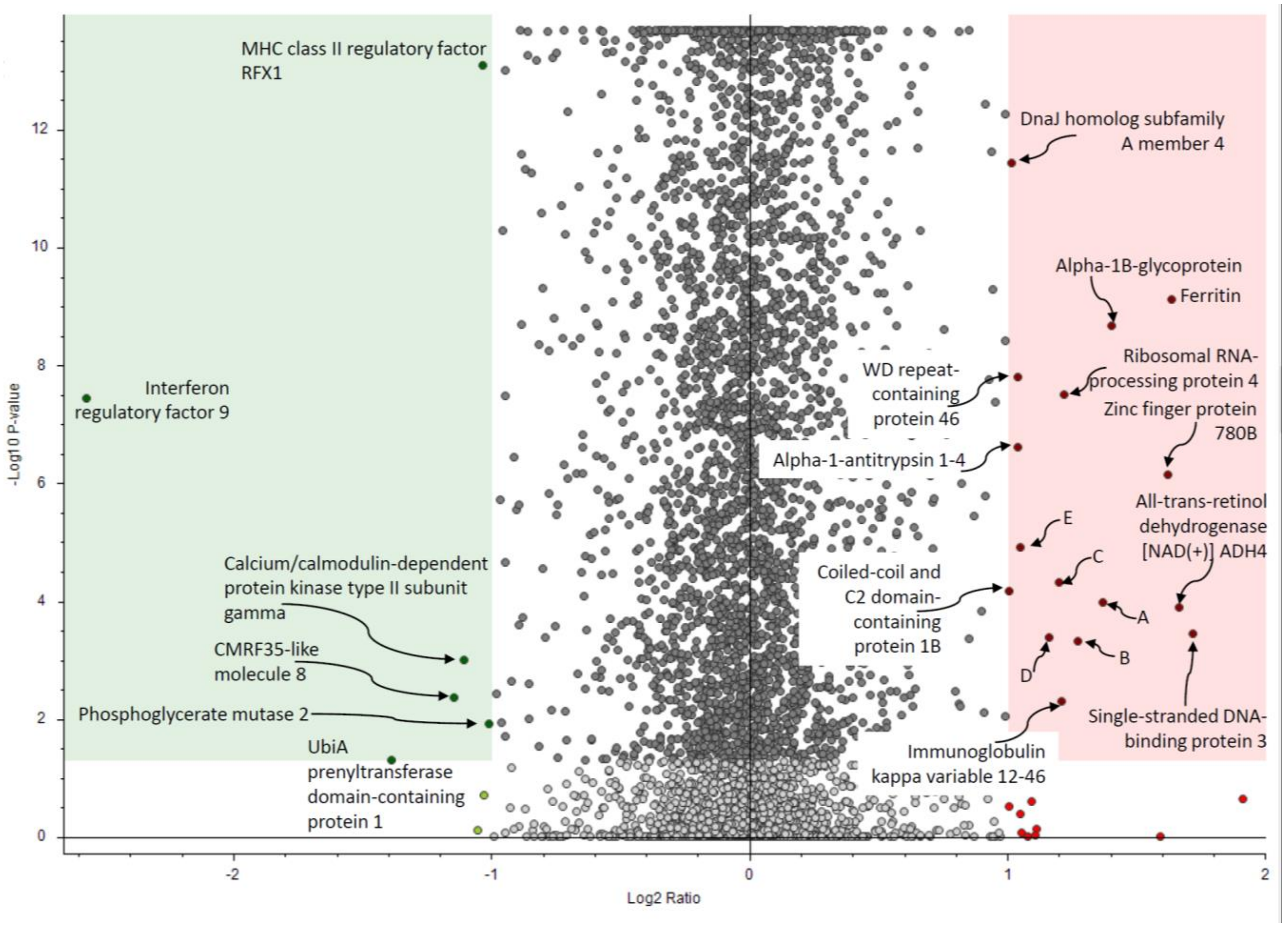


**Figure S5. Male Young vs. Female Young Mass Spectrometry Individual Protein Volcano Plot**

Individually identified proteins were determined to be significantly different if the $\log_2$(ratio of male young vs. female young) was greater than 1 or less than -1 and the $-\log_{10}$(p-value) was greater than 1.25.

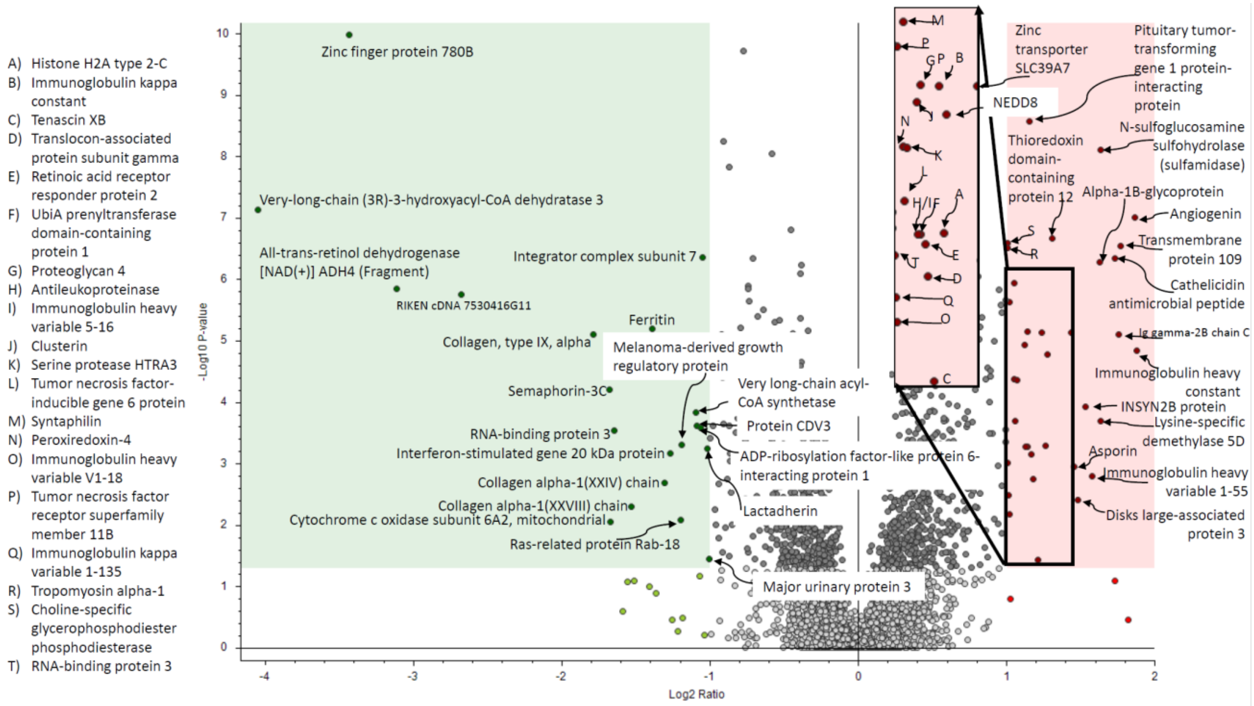


**Figure S6. Male Aged vs. Young Mass Spectrometry Individual Protein Volcano Plot**

Individually identified proteins were determined to be significantly different if the $\log_2$(ratio of male aged vs young) was greater than 1 or less than -1 and the $-\log_{10}$(p-value) was greater than 1.25.

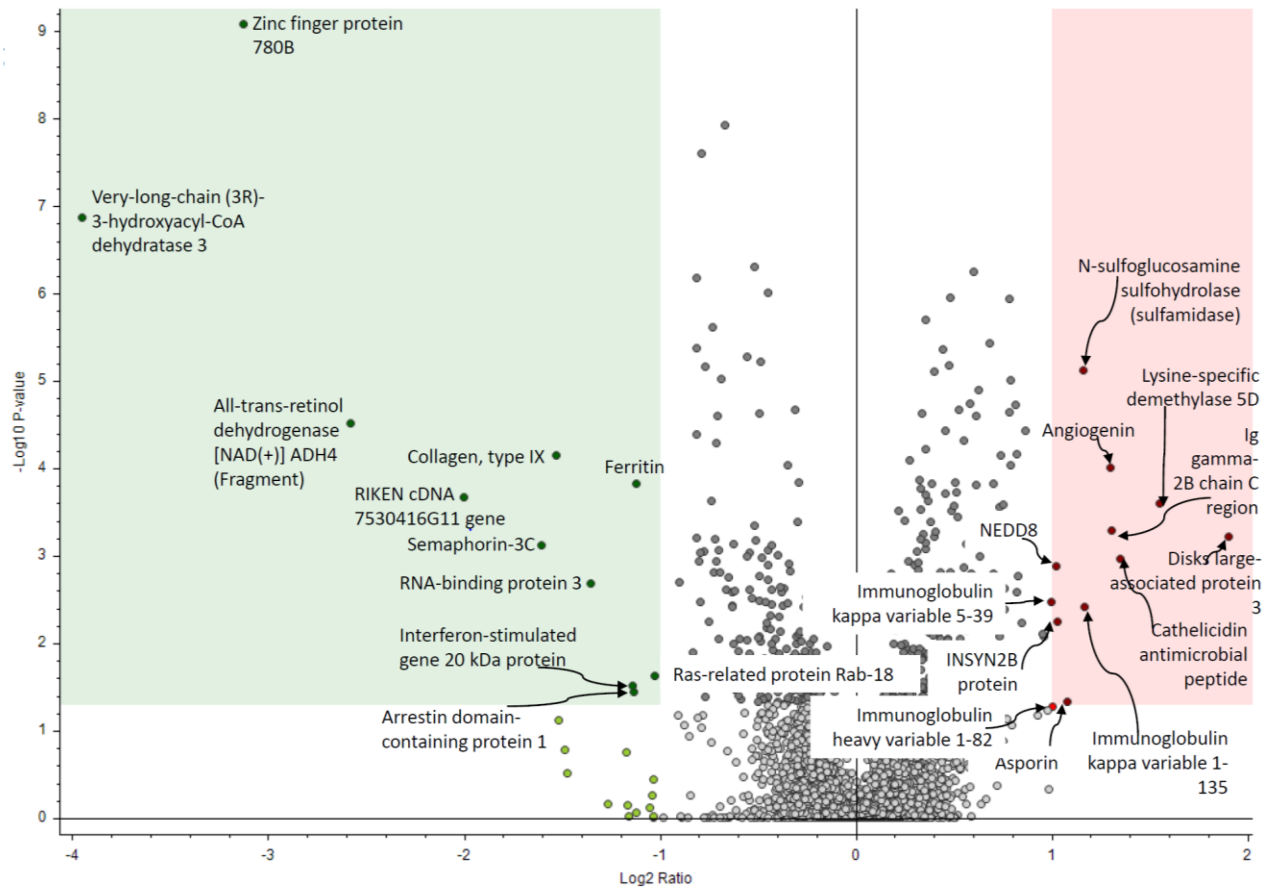


**Figure S7. Male Middle-Aged vs. Young Mass Spectrometry Individual Protein Volcano Plot**

Individually identified proteins were determined to be significantly different if the $\log_2$(ratio of male middle-aged vs young) was greater than 1 or less than -1 and the $-\log_{10}$(p-value) was greater than 1.25.

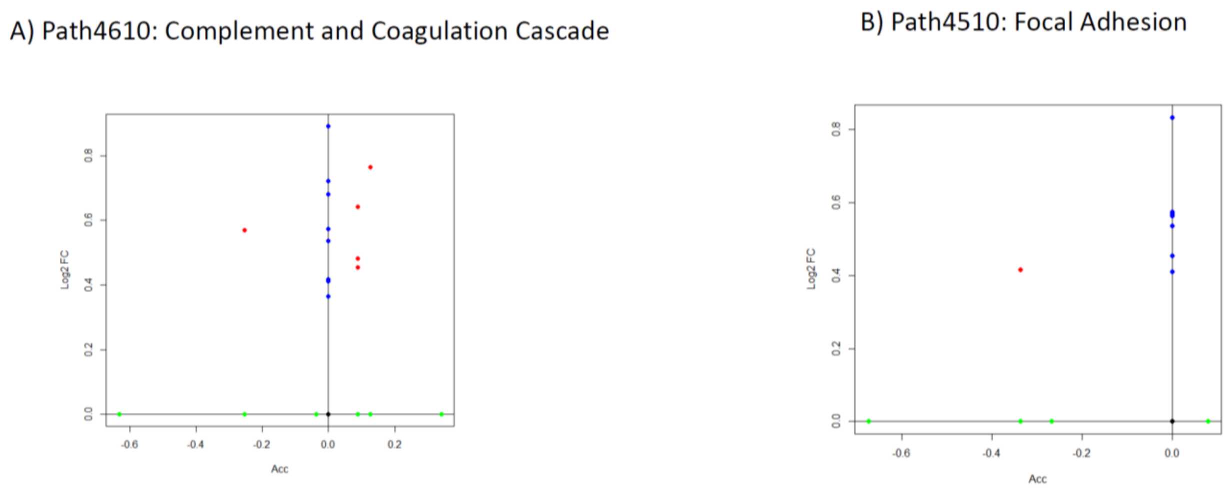


**Figure S8. Female Aged vs. Young ROnToTool Pathway Analysis Statistics**

Individual pathway statistics resulting from Female Aged vs Young Comparison. A) Complement and Coagulation Cascade, B) Focal Adhesion.

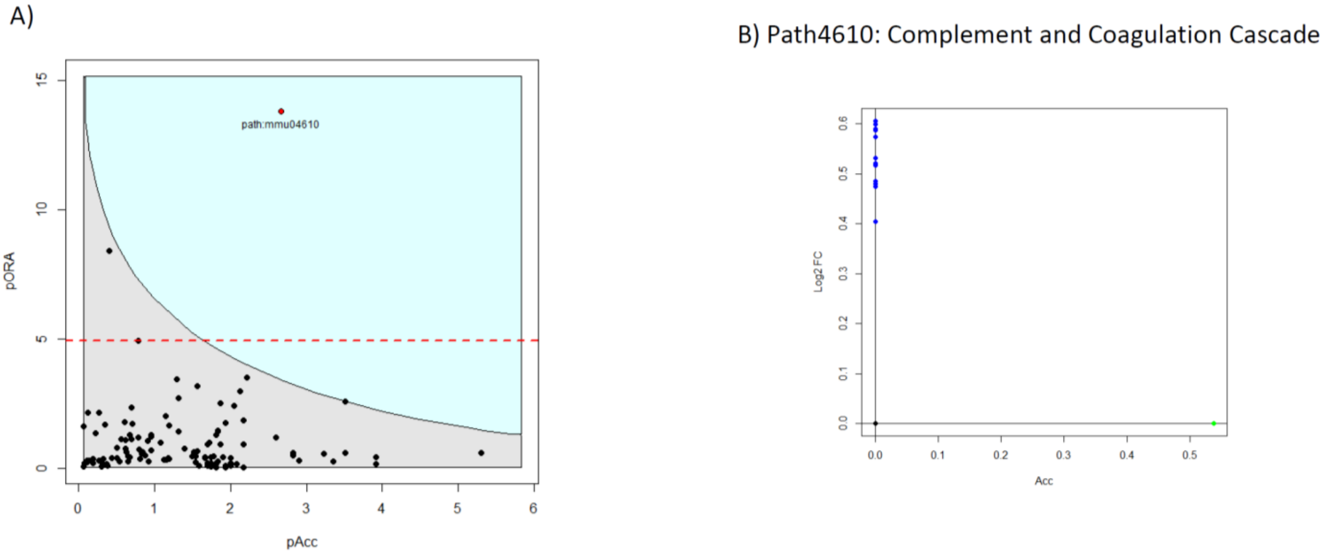


**Figure S9. Female Middle-Aged vs. Young ROnToTool Pathway Analysis Statistics**

Individual pathway statistics resulting from Female Aged vs Young Comparison. A) Whole Pathway Analysis Results, B) Complement and Coagulation Cascade.

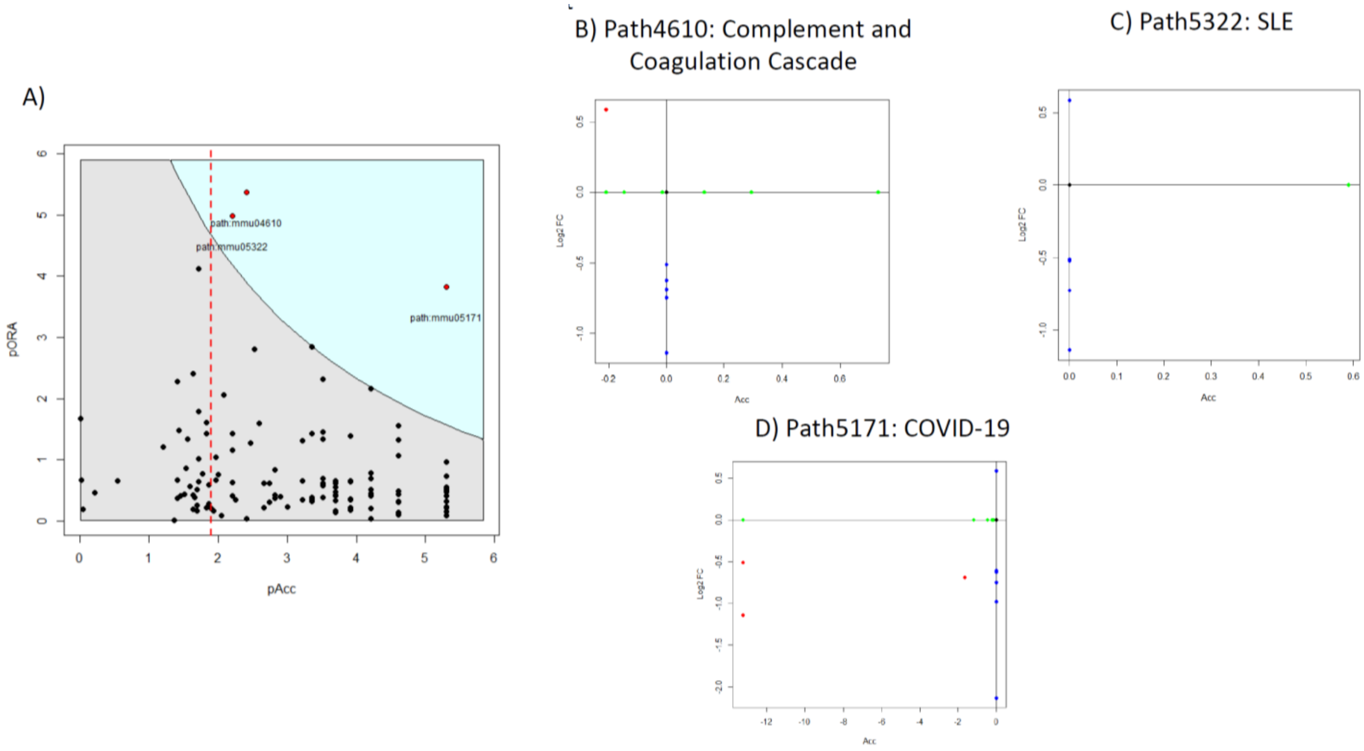


**Figure S10. Male Aged vs. Female Aged ROnToTool Pathway Analysis Statistics**

Individual pathway statistics resulting from Female Aged vs Young Comparison. A) Whole Pathway Analysis Results, B) Complement and Coagulation Cascade, C) SLE, D) COVID-19.

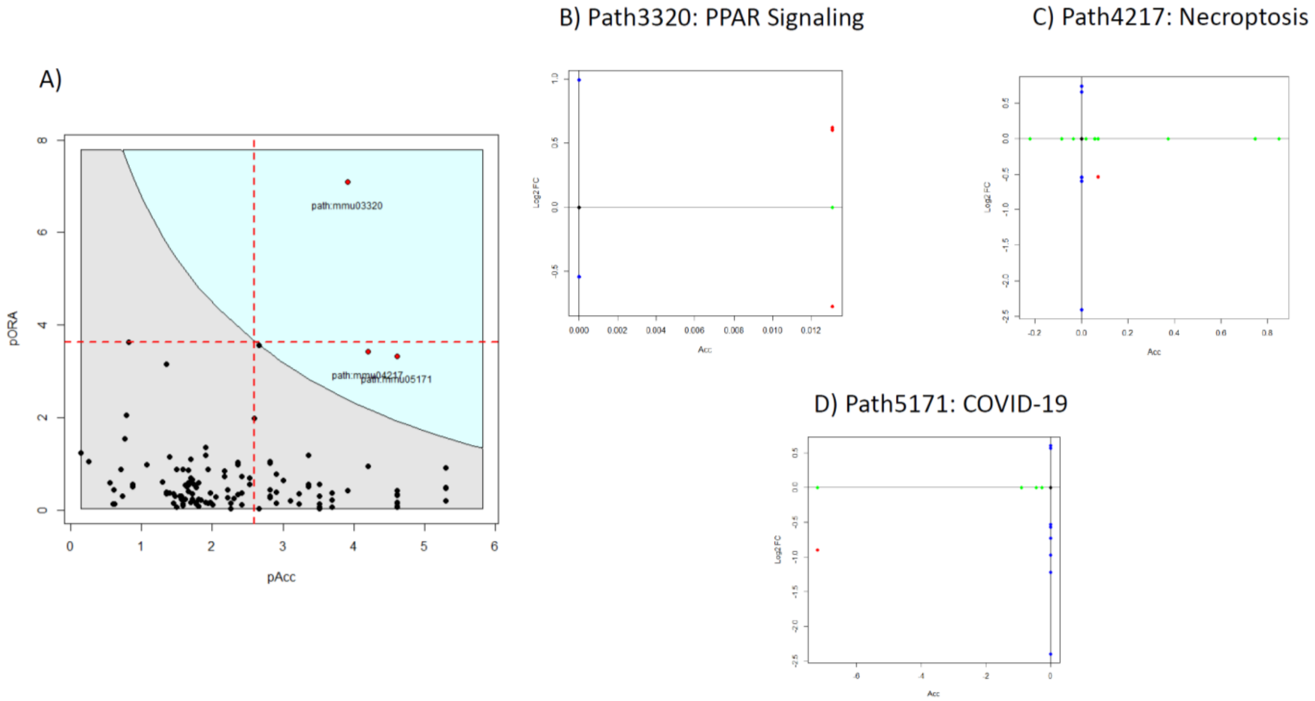


**Figure S11. Male Middle-Aged vs. Female Middle-Aged ROnToTool Pathway Analysis Statistics**

Individual pathway statistics resulting from Female Aged vs Young Comparison. A) Whole Pathway Analysis Results, B) PPAR Signaling, C) Necroptosis, D) COVID-19.

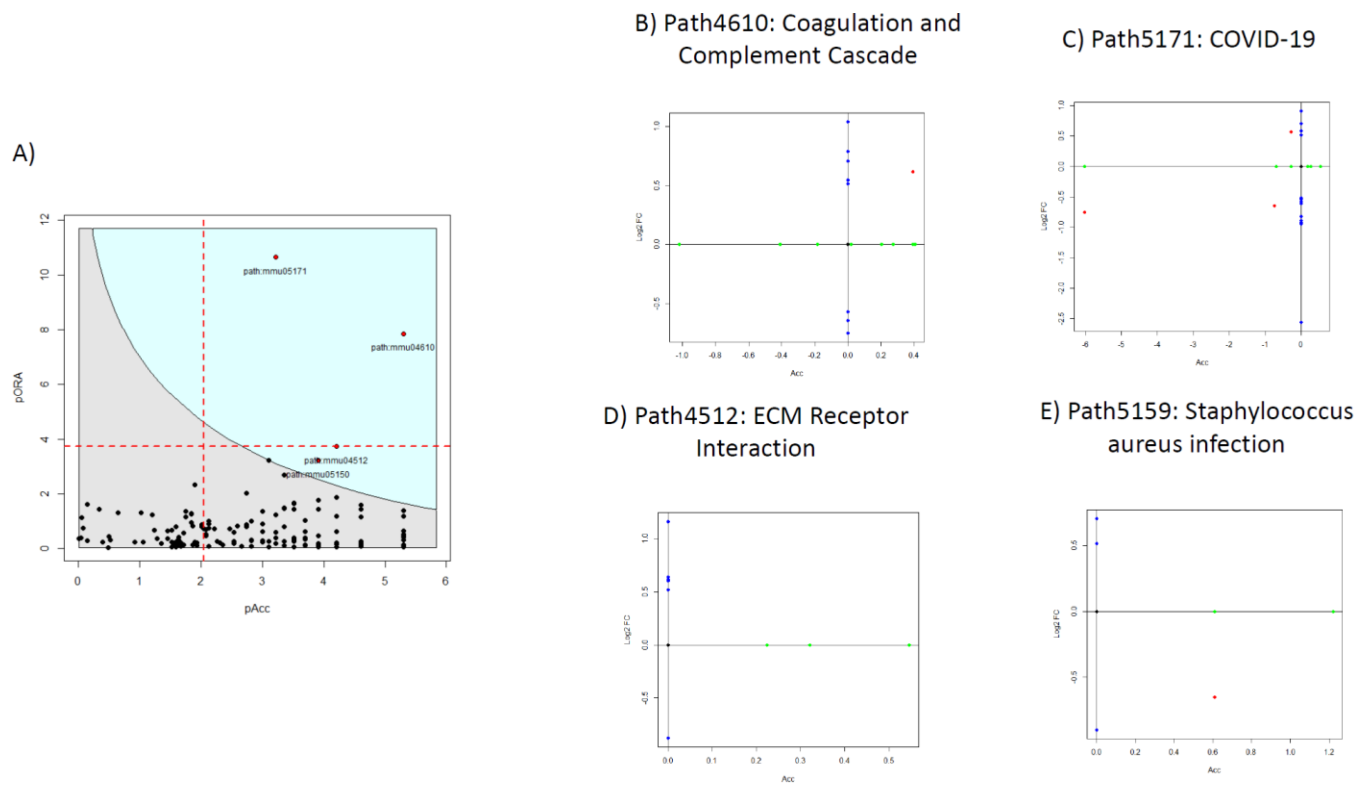


**Figure S12. Male Young vs. Female Young ROnToTool Pathway Analysis Statistics**

Individual pathway statistics resulting from Female Aged vs Young Comparison. A) Whole Pathway Analysis Results, B) Coagulation and Complement Cascade, C) COVID-19, D) ECM Receptor Interaction, E) Staphylococcus aureus infection.

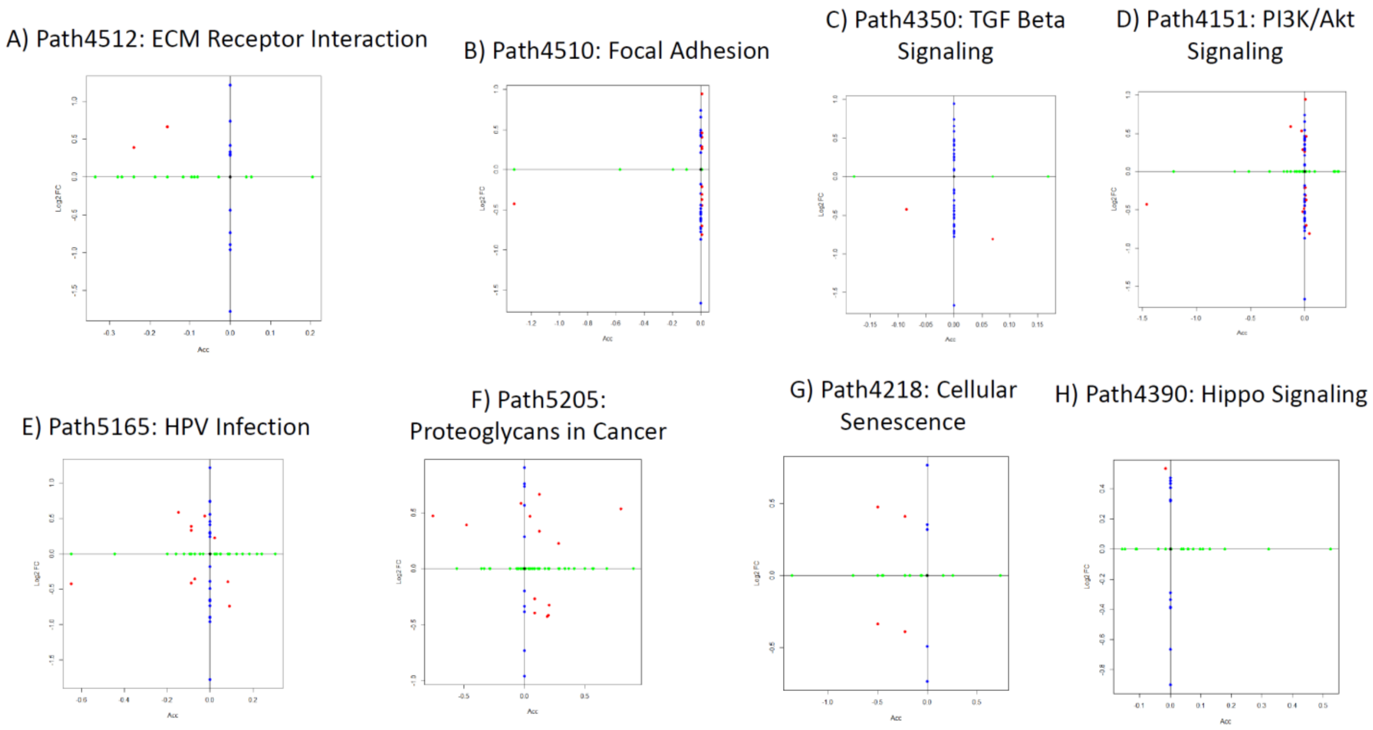


**Figure S13. Male Aged vs. Young ROnToTool Pathway Analysis Statistics**

Individual pathway statistics resulting from Male Aged vs Young Comparison. A) ECM Receptor Interaction, B) Focal Adhesion, C) TGF-Beta Signaling, D) PI3K/Akt Signaling, E) HPV Infection, F) Proteoglycans in Cancer, G) Cellular Senescence, H) Hippo Signaling.

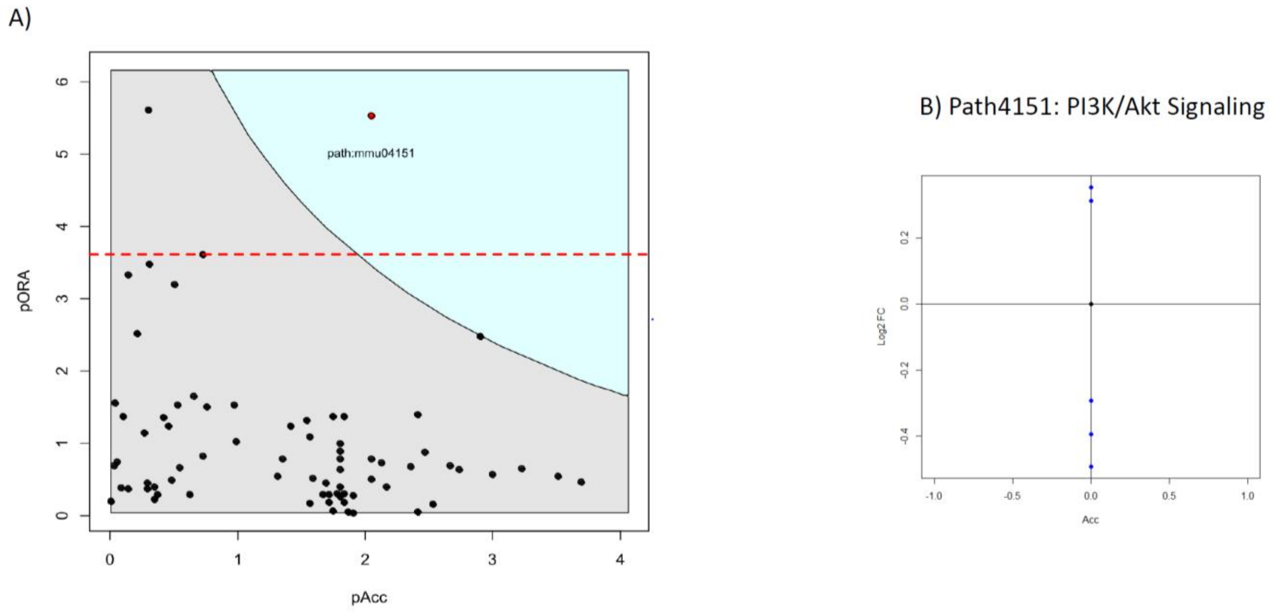


**Figure S14. Male Middle-Aged vs. Young ROnToTool Pathway Analysis Statistics**

Individual pathway statistics resulting from Male Middle-Aged vs Young Comparison. A) Whole Pathways Results, B) PI3K/Akt Signaling.

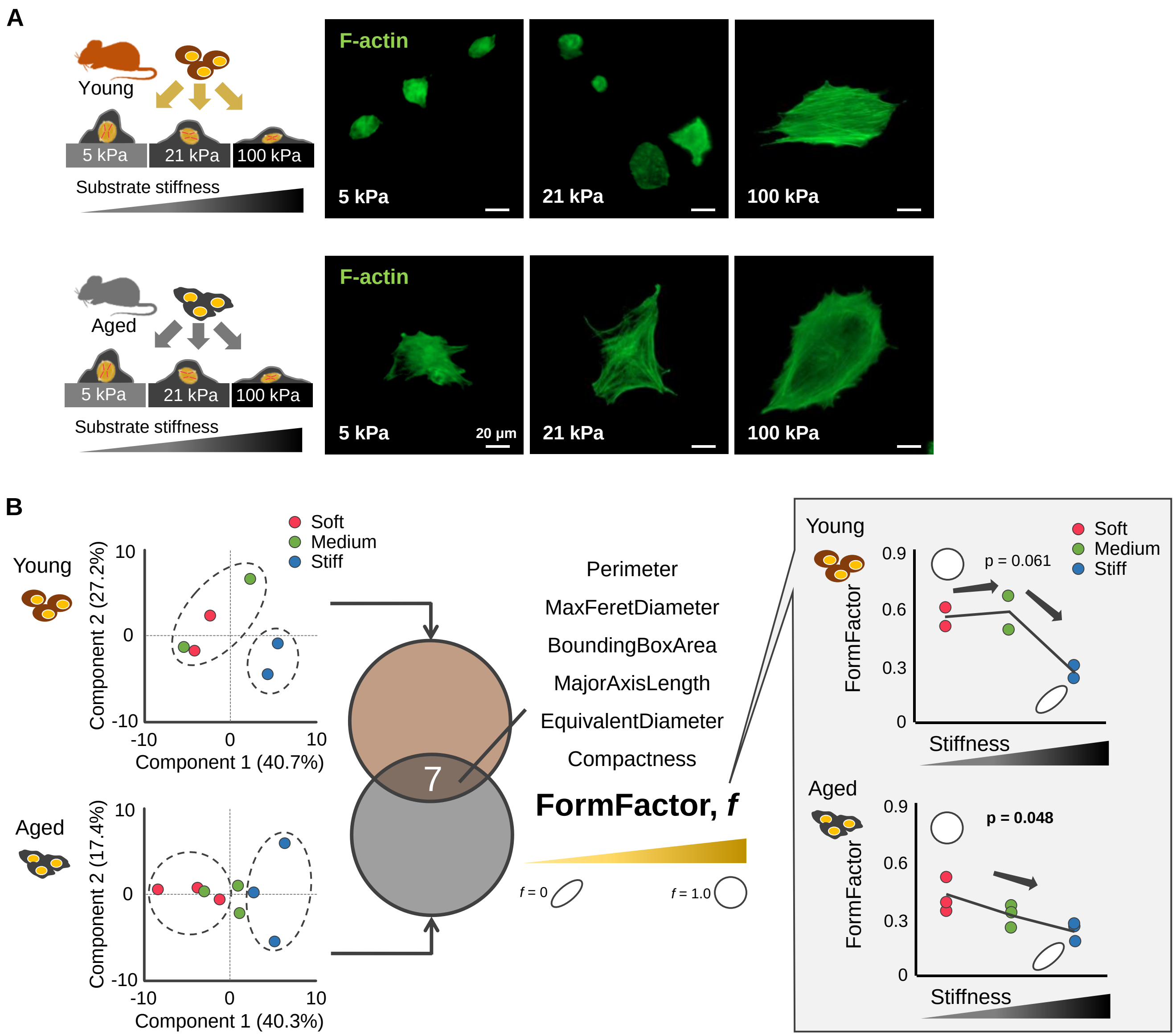


**Figure S15. Chondrocytes morphological changes in different stiffness of polyacrylamide gels**

Primary chondrocytes cultured on physiological range of knee cartilage ECM stiffnesses (5kPa, 21kPa, and 100kPa) displayed different morphology. Representative F-actin image in each stiffness (**A**). Principal component analysis (PCA) of 53 cell morphological variables showed the separate clusters between soft and stiff substrate. Form factor (i.e., roundness) is one of the cell morphological features sensitive to the different stiffness of substrate in both young and aged chondrocytes (**B**).

Table S1. Preparation and characterization of pAAm hydrogels

| pAAm | 40% Acrylamide (μl) | 2% Bis-acrylamide (μl) | 50mM HEPES (ul) (pH 8.2) | TEMED (μl) | 10% Ammonium persulfate (μl) | Substrate stiffness (kPa)* |
|---|---|---|---|---|---|---|
| 5 kPa (soft) | 62.5 | 25 | 412.5 | 1.5 | 5 | 4.9 ± 0.5 |
| 21 kPa (medium) | 150 | 25 | 325 | 1.5 | 5 | 21.2 ± 0.5 |
| 100 kPa (stiff) | 150 | 125 | 225 | 1.5 | 5 | 100.8 ± 2.1 |

*The Young's modulus of pAAm gels was measured by using the atomic force microscopy.

Table S2. Information and dilution protocol of antibodies

| Primary antibodies | Host-species | Product number | Dilution |
|---|---|---|---|
| Type II collagen | Rabbit | Ab34712, Abcam | 1:200 |
| α-Klotho | Rat | MAB1819, R&D systems | 1:100 (tissue), 1:200 (cell) |
| Lamin A/C | Mouse | Sc-376248, Santa Cruz Biotechnology | 1:50 (tissue), 1:100 (cell) |
| Lamin B2 | Rabbit | Ab151735, Abcam | 1:50 (tissue), 1:100 (cell) |
